# MogR is a ubiquitous transcriptional repressor affecting motility, biofilm formation and virulence in *Bacillus thuringiensis*

**DOI:** 10.1101/2020.09.26.311894

**Authors:** Veronika Smith, Malin Josefsen, Toril Lindbäck, Ida K. Hegna, Sarah Finke, Nicolas J. Tourasse, Christina Nielsen-LeRoux, Ole Andreas Økstad, Annette Fagerlund

**Affiliations:** Laboratory for Microbial Dynamics (LaMDa), Section for Pharmacology and Pharmaceutical Biosciences, Department of Pharmacy, University of Oslo, Oslo, Norway; Centre for Integrated Microbial Evolution (CIME), Faculty of Mathematics and Natural Sciences, University of Oslo, Oslo, Norway; Department of Paraclinical Sciences, Faculty of Veterinary Medicine, Norwegian University of Life Sciences, Oslo, Norway; University of Bordeaux, CNRS, INSERM, ARNA, UMR 5320, U1212, Bordeaux, France; Paris-Saclay University, INRAE, AgroParisTech, Micalis, Jouy-en-Josas, France; Nofima, Norwegian Institute of Food, Fisheries and Aquaculture Research, Ås, Norway

**Author notes:** **Correspondence:** Annette Fagerlund, Ole Andreas Økstad.

**Keywords:** *Bacillus cereus* group, MogR, motility regulator, virulence, biofilm

## Abstract

Flagellar motility is considered an important virulence factor in different pathogenic bacteria. In *Listeria monocytogenes* the transcriptional repressor MogR regulates motility in a temperature-dependent manner, directly repressing flagellar- and chemotaxis genes. The only other bacteria known to carry a *mogR* homolog are members of the *Bacillus cereus* group, which includes motile species such as *B. cereus* and *Bacillus thuringiensis* as well as the non-motile species *Bacillus anthracis, Bacillus mycoides* and *Bacillus pseudomycoides*. Furthermore, the main motility locus in *B. cereus* group bacteria, carrying the genes for flagellar synthesis, appears to be more closely related to *L. monocytogenes* than to *Bacillus subtilis,* which belongs to a separate phylogenetic group of Bacilli and does not carry a *mogR* ortholog. Here, we show that in *B. thuringiensis*, MogR overexpression results in non-motile cells devoid of flagella. Global gene expression profiling showed that 110 genes were differentially regulated by MogR overexpression, including flagellar motility genes, but also genes associated with virulence, stress response and biofilm lifestyle. Accordingly, phenotypic assays showed that MogR also affects cytotoxicity and biofilm formation in *B. thuringiensis*. Overexpression of a MogR variant mutated in two amino acids within the putative DNA binding domain restored phenotypes to those of an empty vector control. In accordance, introduction of these mutations resulted in complete loss in MogR binding to its candidate flagellar locus target site *in vitro*. In contrast to *L. monocytogenes*, MogR appears to be regulated in a growth-phase dependent and temperature-independent manner in *B. thuringiensis* 407. Interestingly, *mogR* was found to be conserved also in non-motile *B. cereus* group species such as *B. mycoides* and *B. pseudomycoides*, which both carry major gene deletions in the flagellar motility locus and where in *B. pseudomycoides mogR* is the only gene retained. Furthermore, *mogR* is expressed in non-motile *B. anthracis*. Altogether this provides indications of an expanded set of functions for MogR in *B. cereus* group species, beyond motility regulation. In conclusion, MogR constitutes a novel *B. thuringiensis* pleiotropic transcriptional regulator, acting as a repressor of motility genes, and affecting the expression of a variety of additional genes involved in biofilm formation and virulence.

## 1 Introduction

In many bacterial species, flagella have been demonstrated to be important to virulence functions, including reaching the optimal host site, colonization or invasion, maintenance at the infection site, post-infection dispersal, protein secretion and more (Chaban et al., 2015). Temperature-dependent regulation of motility in the human pathogen *Listeria monocytogenes* has been described in a series of studies published by Higgins and co-workers, in which motility was shown to be regulated by the transcriptional repressor MogR and its anti-repressor GmaR (Gründling et al., 2004; Shen and Higgins, 2006; Shen et al., 2006; Kamp and Higgins, 2009; Shen et al., 2009; Kamp and Higgins, 2011), the genes for which are widely distributed among different *Listeria* spp. (Smith and Hoover, 2009). *L. monocytogenes* is a foodborne facultative intracellular pathogen, which uses flagellum-based motility when present in its extracellular environmental niche. During mammalian infection however, motility genes are downregulated by MogR-dependent repression upon sensing of mammalian physiological temperature (37°C in the human). In this system, the GmaR anti-repressor functions as the temperature sensor, by antagonizing MogR repression activity at temperatures below 37°C (Shen et al., 2006). The activity of GmaR is dependent on the transcriptional activation by DegU at low temperatures, and a temperature-dependent, post-transcriptional mechanism limits GmaR production to temperatures below 37°C (Kamp and Higgins, 2009). This system allows *L. monocytogenes* to switch from an environmental and extracellular motile bacterium to an intracellular pathogen. Inside host cells, flagella are not required as *L. monocytogenes* cells instead move by actin-based motility, and downregulation of flagella during infection is thought to aid bacterial evasion of the host innate immune system (Hayashi et al., 2001; Li et al., 2017).

The only other known bacteria carrying a homolog to *Listeria* spp. *mogR* are species from the *Bacillus cereus* group (*B. cereus sensu lato*) (Gründling et al., 2004; Fagerlund et al., 2010), which is a group of closely related Gram-positive spore-forming bacteria of considerable medical and economic importance. The group comprises at least seven species, including *B. cereus (B. cereus sensu stricto), Bacillus anthracis, Bacillus thuringiensis, Bacillus weihenstephanensis, Bacillus mycoides, Bacillus pseudomycoides,* and *Bacillus cytotoxicus,* which, like *L. monocytogenes,* can be isolated from the environment, e.g. soil, air and water. In contrast to *L. monocytogenes* however, *B. cereus* is an extracellular opportunistic pathogen. The majority of strains of *B. cereus sensu stricto, B. thuringiensis, B. weihenstephanensis* and *B. cytotoxicus* are motile by peritrichous flagella, while *B. anthracis*, *B. mycoides*, and *B. pseudomycoides* are described as non-motile (Twine et al., 2009; Guinebretière et al., 2013). Many strains belonging to the *B. cereus* group can cause foodborne disease (emetic or diarrhoeal syndrome) and serious opportunistic infections in man, but the group also encompasses strains that are used as probiotics (Stenfors Arnesen et al., 2008; Bottone, 2010; Cutting, 2011), some of which have been suggested to have separate species status (e.g. *Bacillus toyonensis;* Jimenez et al., 2013). The highly virulent mammalian pathogen *B. anthracis* is the cause of anthrax disease, being endemic in several parts of the world, and has been used as a biological terror agent. *B. thuringiensis* is an entomopathogenic bacterium, frequently used as a biopesticide to protect crops against insect pests, however this species also carries virulence factors shared with *B. cereus sensu stricto* (Damgaard, 1995; Gaviria Rivera et al., 2000; Swiecicka et al., 2006; Celandroni et al., 2014; Kim et al., 2015) and has caused human infections similar to those caused by *B. cereus* (Samples and Buettner, 1983; Jackson et al., 1995; Damgaard et al., 1997; Hernandez et al., 1998; Ghelardi et al., 2007b). Although strains in the *B. cereus* group to a certain degree follow ecological diversification patterns through evolutionary time (Guinebretière et al., 2008), *B. cereus* and *B. thuringiensis* isolates are intermingled in global MLST- or *k*-mer-based phylogenetic analyses (based on the phylogenetic analysis of common housekeeping genes, or the presence of subsequences of length *k* in a genome, respectively) of a high number of non-biased isolates (Kolstø et al., 2009; Bazinet, 2017), while the thermotolerant species *B. cytotoxicus* forms a separate and phylogenetically remote clade within the *B. cereus* group population.

The bacterial flagellum is a complex molecular structure made up of about 25 different proteins. In most bacteria studied to date, expression of flagellar genes is subject to hierarchical regulation to ensure the sequential expression required for proper flagellum assembly (Smith and Hoover, 2009; Erhardt et al., 2010). In contrast, *L. monocytogenes* and *B. cereus* group strains appear to lack this transcriptional cascade control of flagellar biosynthesis (Smith and Hoover, 2009; Chiara et al., 2015). *B. cereus* group bacteria also lack σ^D^, a common key regulator of motility genes in bacteria, indicating a different mode of transcriptional regulation for the motility genes. In this study, we aimed to investigate functional roles of the MogR homolog identified in *B. cereus* group bacteria, and whether motility regulation in the *B. cereus* group more closely resembles that of *L. monocytogenes* rather than other *Bacillus* species.

## 2 Materials and Methods

### 2.1 Sequence analysis

Motility proteins that were orthologs between species were identified by amino acid sequence searches (BLASTP) performed using BLAST v.2.6.0+ (Altschul et al., 1990; Altschul et al., 1997) between all pairwise combinations of the motility loci from *B. thuringiensis* 407, *L. monocytogenes* EGD-e, and *B. subtilis* 168 (RefSeq accession numbers NC_018877.1, NC_003210.1, and NC_000964.3, respectively; Glaser et al., 2001; Barbe et al., 2009; Sheppard et al., 2013). Results for reciprocal best BLASTP hits between each pair were included in **Supplementary Table 1** if the BLASTP alignments had a percentage of identical matches above 20%, a bit score greater than 30 and covered at least 25% of each amino acid sequence. Comparisons between the motility locus in different *B. cereus* group strains were obtained using the Integrated Microbial Genomes (IMG) browser (Markowitz et al., 2010; http://img.jgi.doe.gov), by searching for orthologous genome neighborhoods to genes from the *B. thuringiensis* 407 motility operon. Annotations for the *B. cereus* ATCC 10987 motility cluster and the *B. mycoides* DSM 2048 *mogR* gene were corrected using EasyGene (Larsen and Krogh, 2003). To perform an exhaustive comparative analysis of the genetic structure of the motility locus, genome sequences of 106 subgroup I strains of the *B. cereus* group that had been sequenced to a minimum of scaffold level were downloaded from NCBI (November 25, 2019), and a local BLAST database was indexed from the corresponding subgroup I proteomes. The corresponding protein sequence from each of the 50 genes in the main *B. thuringiensis* 407 motility locus (genes with locus tags ranging from AFV17362.1 to AFV17411.1, found within coordinates 1608289 to 1653756 in the closed *B. thuringiensis* 407 genome sequence, accession number CP003889) were used as queries in BLASTP searches (parameters: -num_threads 10 -evalue 1.0e-05 -seg no -outfmt 0) for homologous proteins among the subgroup I strains. Output files were parsed using custom-made shell scripts, and *sed/awk*. Candidate MogR binding sites in the *B. thuringiensis* 407 genome were identified by searching the intergenic regions upstream of candidate genes using CLC Main Workbench (Qiagen), with the *L. monocytogenes* consensus MogR binding sequence (TTTTWWNWWAAAA [IUPAC nucleotide codes]; Shen et al., 2009) as query, allowing for up to two mismatches to identify candidate hits.

### 2.2 Strains and growth conditions

The strains used in this study are presented in **Table 1**. *B. thuringiensis* 407 Cry^-^ (also sometimes referred to as Bt407) is an acrystalliferous strain cured of its *cry* plasmid (Lereclus et al., 1989). It is genetically close to the *B. cereus* type strain ATCC 14579 (Tourasse et al., 2006).

**Table 1.** Strains and plasmids used in this study

| Strain or plasmid | Description | Reference or source |
| --- | --- | --- |
| **PLASMIDS** |  |  |
| pHT304-P<sub>xyl</sub> | Low copy number expression/shuttle vector; *xylA* promoter (Ap<sup>r</sup>, Ery<sup>r</sup>) | (Arantes and Lereclus, 1991; Salamitou et al., 1997) |
| pHT304-P<sub>xyl</sub>-*mogR* | wild type *mogR* in pHT304-P<sub>xyl</sub> | This study |
| pHT304-P<sub>xyl</sub>-*mogR*<sup>QN→AA</sup> | Q119A, N120A mutant *mogR* in pHT304-P<sub>xyl</sub> | This study |
| pMAL-p5X | Expression vector for production of MBP fusions, IPTG promoter (Ap<sup>r</sup>) | New England Biolabs |
| pMAL-p5X-*mogR* | wild type *mogR* in pMAL-p5X | This study |
| pMAL-p5X-*mogR*<sup>QN→AA</sup> | Q119A, N120A mutant *mogR* in pMAL-p5X | This study |
| **STRAINS** |  |  |
| ***B. thuringiensis*** |  |  |
| 407 | *B. thuringiensis* 407 Cry<sup>-</sup> | (Lereclus et al., 1989) |
| 407/pHT304-P<sub>xyl</sub> | pHT304-P<sub>xyl</sub> in 407 | This study |
| 407/MogR<sup>+</sup> | pHT304-P<sub>xyl</sub>-*mogR* in 407 | This study |
| 407/MogR<sup>QN→AA</sup> | pHT304-P<sub>xyl</sub>-*mogR*<sup>QN→AA</sup> in 407 | This study |
| 407Δ*flaAB* | *B. thuringiensis* 407Δ*flaA*Δ*flaB* (Km<sup>r</sup>) | (Houry et al., 2010) |
| 407Δ*flaAB*/pHT304-P<sub>xyl</sub> | pHT304-P<sub>xyl</sub> in 407Δ*flaAB* | This study |
| 407Δ*flaAB*/MogR<sup>+</sup> | pHT304-P<sub>xyl</sub>-*mogR* in 407Δ*flaAB* | This study |
| 407Δ*flaAB*/MogR<sup>QN→AA</sup> | pHT304-P<sub>xyl</sub>-*mogR*<sup>QN→AA</sup> in 407Δ*flaAB* | This study |
| ***E. coli*** |  |  |
| BL21(DE3) | *E. coli* BL21(DE3) | New England Biolabs |
| BL21/MogR<sup>+</sup> | pMAL-p5X-*mogR* in BL21(DE3) | This study |
| BL21/MogR<sup>QN→AA</sup> | pMAL-p5X-*mogR*<sup>QN→AA</sup> in BL21(DE3) | This study |

Unless otherwise stated, *B. thuringiensis* 407 cultures were inoculated with 1% of an overnight culture and grown at 30°C and 200 rpm in Luria Bertani (LB) broth or in bactopeptone medium (1% w/v bactopeptone, 0.5% w/v yeast extract, 1% w/v NaCl). For cloning and expression in *Escherichia coli,* ampicillin at 50 or 100 μg mL^−1^, kanamycin at 50 μg mL^−1^ and/or erythromycin at 400 μg mL^−1^ was used. Erythromycin at 10 μg mL^−1^ was used to maintain the pHT304-P_*xyl*_ plasmid constructs in *B. thuringiensis* 407. For induction of gene expression from the *xylA* promoter on pHT304-P_*xyl*_, xylose was added to the growth medium at 1 mM or as otherwise stated. For induction of MBP-MogR in *E. coli* BL21(DE3), 0.3 mM isopropyl-β-D-thiogalactopyranoside (IPTG) was used. Growth curves were prepared in bactopeptone medium, pH 7.0 with cultures grown with shaking at 220 rpm, using 50 mL culture volumes in 250 mL baffled flasks.

### 2.3 Reverse transcription quantitative PCR (RT-qPCR)

For analysis of *mogR, flaA* and *flaB* expression throughout the bacterial growth phase, RT-qPCR was performed essentially as described by Fagerlund et al. (2016). Briefly, cells grown in bactopeptone medium at 30°C were incubated in an equal volume of ice-cold methanol for 5 min before harvesting by centrifugation. Cells were lysed using a Precellys 24 Tissue Homogenizer (Bertin) and RNA was isolated using the RNeasy Mini or Midi Kits (Qiagen). After treatment with DNase and further purification, cDNA synthesis was performed in duplicate for each sample using SuperScript III Reverse Transcriptase (Invitrogen). For all samples, a negative control reaction without reverse transcriptase was included. RT-qPCR was carried out with a LightCycler 480 Real-Time PCR System (Roche) using primers listed in **Supplementary Table 2**. The three genes *gatB*, *rpsU*, and *udp*, shown to be stably expressed throughout the *B. cereus* life cycle (Reiter et al., 2011), were used as reference genes, and were included for each sample and on each plate. The second derivative maximum method in the LightCycler 480 software (Roche) was utilized to obtain a quantification cycle (Cq) value for each reaction. The expression of each target gene in each biological replicate was converted into E^Cq^ values (Pfaffl, 2001) and then normalized to the geometric mean of the E^Cq^ values obtained for the three reference genes. Finally, averages and standard deviations were calculated from the normalized expression values from the biological replicates.

For analysis of *flaA* and *flaB* expression in the *B. thuringiensis* 407 strains overexpressing MogR and MogR^QN→AA^ (see below), the same procedure was used except that *gatB* and *rpsU* were used as reference genes.

### 2.4 Cloning of *mogR* for expression in *B. thuringiensis* 407

The low-copy number *E. coli/Bacillus* shuttle vector pHT304-P_*xyl*_, in which *xylR* and the *xylA* promoter from *B. subtilis* was inserted into the pHT304 cloning site (Arantes and Lereclus, 1991) allowing xylose-inducible expression of downstream cloned genes, was used for overexpression studies. The sequence encoding MogR from *B. thuringiensis* 407 (BTB_RS08390) was PCR-amplified using primers 5’ -gtc<u>ggatcc</u>gaattgtgaaaggatgagg-3’ and 5’-taa<u>ggtacc</u>tcttccttcttcggaacg-3’ and genomic DNA as the template. The PCR product was inserted into pHT304-P_*xyl*_ using the primer-incorporated *Bam*HI and *Kpn*I restriction sites (underlined), creating pHT304-P*_xyl_-mogR* and placing *mogR* under transcriptional control of the *xylA* promoter.

The plasmid pHT304-P*_xyl_-mogR^QN→AA^*, containing *mogR* with amino acid substitutions in two of the six conserved amino acid residues predicted to make base-specific contacts with the MogR recognition site (Shen et al., 2009) was created by site-directed mutagenesis of *pHT304-P_xyl_-mogR* using the QuikChange II Site-Directed Mutagenesis Kit (Stratagene) and primers 5’-tccaaaaacagaaagtcaattg<u>gc</u>a<u>gc</u>tacgtattataaattgaaaaaacgtg-3’ and 5’-cacgttttttcaatttataatacgta<u>gc</u>t<u>gc</u>caattgactttctgtttttgga-3’ (mutated bases underlined). The introduced mutations were Q119A and N120A.

The plasmids were verified by sequencing. The constructed pHT304-P_*xyl*_ plasmids and the empty vector were introduced by electroporation into *B. thuringiensis* 407 (Masson et al., 1989).

### 2.5 Motility assay

Swimming ability was determined on 0.3% LB soft agar plates with 1 mM xylose and 10 μg mL^−1^ erythromycin added. A 5 μL drop of culture grown in LB broth overnight at 30°C was spotted on each agar plate. The plates were wrapped in plastic and incubated for 7 h at 30°C. Each independent assay was performed with two or three technical replicates.

### 2.6 Atomic force microscopy (AFM)

AFM imaging and analysis was performed using a Nanowizard AFM microscope (JPK Instruments). Bacterial cell culture was grown in LB broth with erythromycin (10 μg mL^−1^) and xylose (10 mM) at 37°C to an OD_600_ of 3. One mL of culture was washed three times and finally resuspended in 0.9% NaCl. Cells were then diluted (15:50) in a 10 mM magnesium/Tris buffer, pH 7.5. Ten μL was applied onto freshly cleaved mica surfaces mounted on a glass slide, and allowed to adhere for 10 min followed by washing (10×100 μL) using sterile dH_2_O. Excess water was carefully removed, and the slide gently dried using a nitrogen gas jet stream. Images were recorded in intermittent-contact mode at room temperature in air using a MicroMasch NSC35/AIBS cantilever. AFM images were analyzed using The NanoWizard IP Image Processing Software (JPK Instruments).

### 2.7 SDS-PAGE and Western immunoblotting

For detection of flagellin, for each strain analyzed, two parallel bacterial culture samples (10 mL LB broth) were harvested by centrifugation (4100 × *g*, 4°C) after 3.5 h growth at 30°C (OD_600_ ~1.2) – one for extraction of surface proteins, and one for whole cell protein extraction. The cell pellets were resuspended in 1 mL PBS (pH 7.1) and kept on ice. For extraction of surface proteins, the washed cells were centrifuged for 5 min at 16,000 × *g* and 4°C and resuspended in an equal volume of 2×SDS-PAGE sample buffer before incubation at 95°C for 5 min. The supernatant was collected by centrifugation as before. The whole cell fraction was prepared by washing the cell pellet in PBS followed by centrifugation for 5 min at 16,000 × *g* and 4°C. The pellet was then resuspended in 500 μL PBS and lysed using a Precellys 24 Tissue Homogenizer (Bertin). The supernatant was collected by centrifugation for 8 min as before. Whole cell supernatant (21 μL) with 7 μL 4×SDS-PAGE buffer and 5 μL of the surface protein fraction were separated on 12% SDS-PAGE gels as described below.

For detection of Hbl, Nhe and CytK, cultures were harvested after 4.5 h growth. Supernatant samples were collected by centrifugation, concentrated 40-fold by precipitation with four volumes of ice-cold acid acetone:methanol (1:1 v/v), stored at −20°C overnight, and harvested by centrifugation for 30 min at 16,000 × *g* and 4°C. Then, pellets were left to evaporate at 4°C overnight and resuspended in 2×SDS-PAGE sample buffer. Samples were diluted 20-fold with MQ H_2_O. Concentrated supernatant samples (10 μL) were separated on 10% SDS-PAGE gels.

SDS-PAGE was carried out using a Bio-Rad Mini-Protean II Dual Slab Cell, using 10 μL Prestained Protein Marker, Broad range (New England Biolabs) as the molecular weight marker. Western blot analysis was performed using Immun-Blot PVDF membranes (Bio-Rad) according to standard protocols (Harlow, 1988). Blocking was performed for 1 h in 5% non-fat dry milk in TBST. Flagellin proteins were detected using a rabbit antiserum raised against flagellin from *Bacillus mojavensis,* used at a 1:300 dilution, and a HRP-conjugated donkey-anti-rabbit antibody (Santa Cruz Biotechnology) diluted 1:10,000 as secondary antibody. CytK was detected using rabbit antiserum (Fagerlund et al., 2004) diluted 1:2000, followed by an HRP-conjugated donkey anti-rabbit antibody (Santa Cruz Biotechnology) diluted 1:5000. Hbl B and NheA were detected using monoclonal antibodies 2A3 against Hbl B (Dietrich et al., 1999) and 1A8 against NheA (Dietrich et al., 2005) (both diluted 1:15), followed by HRP-conjugated AffiniPure Goat-anti-mouse IgG (H+L) (Jackson Immuno Research Laboratories) at 80 μg mL^−1^. SuperSignal West Femto Substrate (Pierce) was used to develop the blots. Western blots were photographed in a Chemi Genius Bio Imaging System (Syngene), and sub-saturation images were saved as jpeg files in ImageJ (Abràmoff et al., 2004).

### 2.8 Microarray analysis

Cultures of *B. thuringiensis* 407 harboring either pHT304-P*_xyl_* or pHT304-P*_xyl_-mogR* were grown in LB broth containing 10 mM xylose at 37°C for 3 h, and then incubated in an equal volume of ice-cold methanol for 5 min before pellets were harvested by centrifugation at 2800 × *g* for 20 min. For extraction of RNA, cells were lysed using a Precellys 24 Tissue Homogenizer (Bertin) and RNA was isolated using the RNeasy Mini Kit (Qiagen) and subjected to on-column DNase-treatment using the RNase-Free DNase Set (Qiagen). cDNA synthesis, labelling and purification was performed as described (Gohar et al., 2008). Microarray slides were printed at The Microarray core facility of the Norwegian University of Science and Technology (NTNU). Design, printing, prehybridization, hybridization and scanning of the slides and analysis of the data was performed as described (Gohar et al., 2008). Each microarray experiment was based on four slides, all biological replicates. *P*-values were computed using a false discovery rate (FDR) of 0.05.

The microarray slides contain 70-mer oligonucleotide probes designed to detect open reading frames (ORFs) in the following strains: *B. anthracis* Ames, *B. anthracis* A2012, and *B. cereus* ATCC 14579, in addition to selected genes from *B. cereus* ATCC 10987 (Kristoffersen et al., 2007). All probe sequences on the microarray were analyzed by BLAST for hits to the annotated genes of a *B. thuringiensis* 407 draft genome sequence (as of April 30, 2009; the gene lists are based on the GenBank annotations as of this date; accession no. ACMZ00000000.1). Only probes with 93% identity or greater to a transcript/feature sequence of *B. thuringiensis* 407 were included in the analysis. Of the predicted *B. thuringiensis* 407 genes, 1719 genes, most of which were hypothetical genes, did not have corresponding probes on the array (these include the 761 genes on contigs 00213 and 00060, which appear to be plasmid-borne). COG categories were obtained for the analysed genes as reported in the IMG database (http://img.jgi.doe.gov).

### 2.9 Cytotoxicity assay

The Vero cell cytotoxicity assay was performed as described by Lindbäck and Granum (2006), and measures the percentage inhibition of C^14^-leucine incorporation in cells due to the cells being subjected to toxins, calculated relative to a negative control where cells were not subjected to toxin sample. Samples of early stationary phase cultures grown to an OD_600_ of 2.4 were collected and 100 μL or 150 μL samples (for deletion mutants and overexpression strains, respectively) were applied to the cytotoxicity assay. The assays were performed for three independent biological replicates with two technical replicates in each assay.

### 2.10 Virulence in *Galleria mellonella* larvae

The virulence-related properties of MogR were assessed by comparing the killing effect of the *B. thuringiensis* 407 MogR overexpression and empty vector control (pHT304-P*_xyl_* plasmid) strains, in both wild type and Δ*flaAB* backgrounds, by infection (force feeding) in 5^th^ instar *Galleria mellonella* larvae. *G. mellonella* eggs were hatched at 28°C and the larvae reared on beeswax and pollen. For infection experiments, groups of 20 to 25 *G. mellonella* larvae, weighing about 200 mg each, were used. As MogR overexpression is activated from the pHT304-P*_xyl_* promoter by the addition of xylose, we tried for the first time to evaluate if activation could occur in the insect intestine as well. Therefore, xylose (20 mM) was both added to the LB growth medium of the four strains, as well as to the bacterial inoculums and the toxin alone control (Cry1C) at time zero (time point of force feeding). Larvae were force fed a second time with 10 μL 20 mM xylose 5 h later (in order to again activate MogR expression from the plasmid). Infections were otherwise performed as previously described (Fedhila et al., 2006), by force feeding with 10 μL of a mixture containing 4-5×10^6^ of vegetative bacteria (exponential growth OD_600_ ~1 in LB) with 20 mM xylose and 3 μg of activated Cry1C toxin to overcome the *B. thurinngiensis* 407 strain being *cry* negative. The larvae in the control group were fed PBS buffer or Cry1C toxin and xylose in corresponding amounts to the samples containing bacterial inocula. The chosen dose was expected to result in about 70% (± 5%) mortality at 37°C after 48 h.

### 2.11 Biofilm assays

The ability to form biofilms was determined using a glass tube screening assay (Houry et al., 2010). Briefly, exponential phase cultures were diluted into HCT medium (Lecadet et al., 1980; Lereclus et al., 1982) to an OD_600_ of 0.01, and 2 mL was inoculated into sterile 6 mL glass tubes. The tubes were incubated for 72 h at 37°C. The biofilm was subsequently collected by removing the culture medium with a Pasteur pipette and thoroughly vortexing in 2 mL PBS before measuring the OD_600_ of the suspension of biofilm cells. Each strain was tested in five biological replicates, each with 3 or 4 technical replicates.

The ability to form biofilms in polyvinylchloride (PVC) microtiter plates was determined using a crystal violet biofilm screening assay (Auger et al., 2006). Briefly, fresh bactopeptone medium was inoculated with 0.5% exponential phase culture, transferred to 96-well plates (Falcon #353911) and incubated for 24, 48 and 72 h at 30°C. The biofilm was subsequently washed using PBS, stained using 0.3% crystal violet, washed as before, solubilized with acetone:methanol (1:3 v/v), and transferred to flat-bottomed microtiter plates (Falcon #353915) for determination of the absorbance of the solubilized dye at 570 nm. Each strain was tested in triplicate. Statistical analyses were performed separately for each time point, as described under “Statistical analyses” below.

### 2.12 Expression of MBP-tagged MogR in *E. coli*

The sequence encoding MogR from *B. thuringiensis* 407 (BTB_RS08390) was PCR amplified using primers 5’-atgtatcaccacacagcaattaatgtattag-3’ and 5’-gcgc<u>ggatcc</u>ttattactgtgttacggtcataacttgtcc-3’ and genomic DNA as the template. The PCR product was cloned into the pMAL-p5x vector (New England Biolabs) using the *Xmn*I and *Bam*HI restriction sites (underlined) according to the manufacturer’s instructions, allowing expression of MBP-MogR. A construct expressing the MBP-MogR^QN→AA^ protein was created by site-directed mutagenesis of pMAL-p5x-*mogR* using the QuikChange II Site-Directed Mutagenesis Kit (Stratagene) with the same primers as for pHT304-P*_xyl_*-MogR^QN→AA^. Constructs were transformed into *E. coli* BL21(DE3) cells. The plasmids were verified by sequencing. MBP-MogR and MBP-MogR^QN→AA^ proteins were expressed in *E. coli* BL21(DE3) and purified using the manufacturer’s manual “Method I – Total cell extract”. In short, 100 mL LB broth was inoculated with a 1 mL overnight culture containing cells harbouring fusion plasmid. Cells were grown at 37°C with shaking to an OD_600_ of ~0.5. IPTG was added to a final concentration of 0.3 mM and cells were induced for 2 h. The cells were harvested by centrifugation at 4500 × *g* for 10 min and frozen at −20°C overnight, sonicated in an ice-water bath, and then centrifuged at 20,000 × *g* for 20 min at 4°C. The supernatants were diluted 1:6 in Column Buffer (20 mM Tris-HCl, 200 mM NaCl, 1 mM EDTA). MBP-MogR proteins were purified by amylose affinity chromatography (Poly-Prep Chromatography column, 0.8 × 4 cm, Bio-Rad). The identity and purity of proteins was confirmed by SDS-PAGE (Coomassie stain) and the quantity determined by Bradford assay (Pierce BCA Protein Assay Kit, Thermo Fisher).

### 2.13 Electrophoretic mobility shift assay (EMSA)

DNA fragments were generated by PCR using chromosomal DNA from *B. thuringiensis* 407 as template and the following primer combinations: 5’-cgggtgtcaactaaaaattcg-3’ / 5’-caactatcataatatcaccttttcgg-3’ (*fla),* 5’-ttacagaaatgaaatttacggataac-3’ / 5’-ccttatcctttctgtctggtc-3’ (*hbl),* and 5’-tccgtatgtaattccgtttcaaga-3’ / 5’-aatttcctgcttgacccctt-3’ (ctrl1). Corresponding 5’ biotinylated probes were generated for the *fla* and *hbl* fragments using biotin-labelled primers. The ctrl1 fragment was chosen as a nonspecific competitor control for binding to *B. thuringiensis* DNA and is a PCR product from inside a randomly chosen gene from *B. thuringiensis* 407 (BTB_RS08990 encoding an elongation factor G-binding protein). A commercial EMSA kit (LightShift Chemiluminescent, Thermo Scientific) was used to detect binding of MogR protein to DNA following the manufacturer’s instructions. Unlabelled Epstein Barr nuclear antigen DNA supplied with the kit was used as non-*Bacillus* nonspecific competitor DNA control (ctrl2 fragment). DNA binding reactions were conducted in 20 μL volumes containing 10 mM Tris-HCl pH 7.5, 50 mM KCl, 1 mM DTT, and 50 ng μL^−1^ polydeoxyinosinicdeoxycytidylic acid (poly dI-dC). In each reaction, MBP-MogR or MBP-MogR^QN→AA^ was added to approximately 2 fmol biotinylated DNA probe and incubated for 20 min at room temperature. Each binding reaction was loaded on a 5% SDS-PAGE gel (0.5×TBE) and resolved for 1 h at 100 V. Gels were blotted onto a nylon membrane and DNA was crosslinked to the membrane by exposure to UV light (120 mJ/cm^2^) for 60 seconds. Finally, biotin-labelled DNA was detected using chemiluminescense as described in the EMSA kit instructions.

### 2.14 Quantification of cyclic di-GMP by LC-MS/MS

LC-MS/MS analysis was based on a method described by Spangler et al. (2010), with some modifications, as previously described (Fagerlund et al., 2016), using a Thermo Scientific LTQ XL Linear Ion Trap Mass Spectrometer (Thermo Scientific) and separation on a 5 mm × 1 mm I.D. Nucleodur C18 Pyramid precolumn and a 50 mm × 1 mm I.D. Nucleodur C18 pyramid analytical column (both from Marchery-Nagel), with an electrospray ionization (ESI) source operated in the positive ionization mode to interface the HPLC and the MS.

### 2.15 Statistical analyses

Minitab v.17 software was used for statistical analysis. Unless stated otherwise, all data were analyzed using analysis of variance (ANOVA) followed by Tukey’s *post hoc* test for pairwise comparisons. *P*-values <0.05 were regarded as significant.

## 3 Results

### 3.1 The motility gene loci in *B. thuringiensis* and *L. monocytogenes* are closely related

A cluster of approximately 45-50 genes with homology to flagellar-based motility- and chemotaxis genes is present in most *B. cereus* group strains. Although *B. cereus* group bacteria are phylogenetically more closely related to *B. subtilis* than to *L. monocytogenes*, some aspects of organization of the *B. thuringiensis* 407 flagellar motility gene cluster were found to more closely resemble that of *L. monocytogenes*. Bidirectional BLAST analysis showed that a greater number of motility protein orthologs is shared between *B. thuringiensis* 407 and *L. monocytogenes* EGD-e (41 orthologous proteins, average alignment percentage identity 46%) than between *B. thuringiensis* 407 and *B. subtilis* 168 (31 orthologous proteins, 37% average sequence identity) (**Supplementary Table 1**). Furthermore, only other *Listeria* spp. and species of the *B. cereus* group are known to have orthologs to the *L. monocytogenes* <u>mo</u>tility <u>g</u>ene <u>r</u>epressor MogR (Gründling et al., 2004; Smith and Hoover, 2009), potentially indicating that elements of motility regulation may be shared between these species (**Figure 1** and **Supplementary Table 1**).

**Figure 1.**
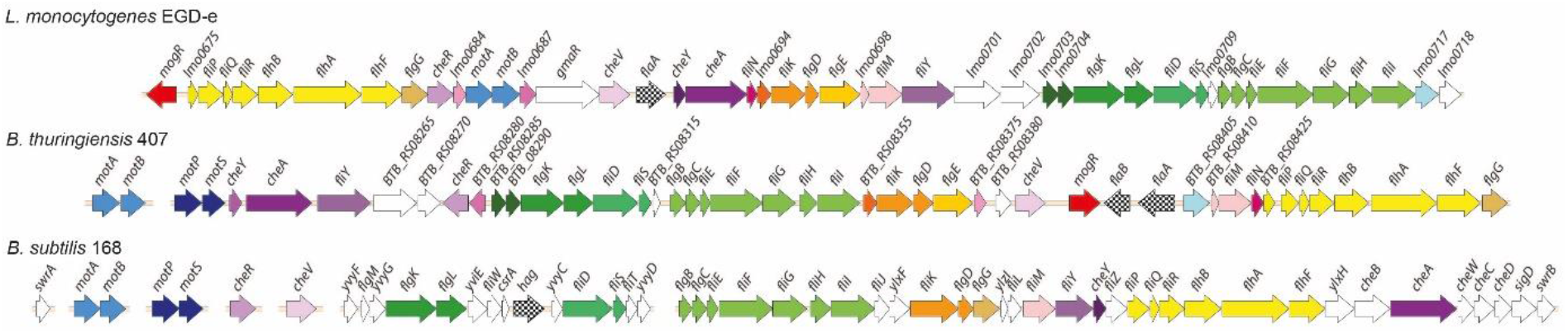
Comparison of genomic loci encoding flagellar-based motility genes in *Listeria monocytogenes, Bacillus thuringiensis* and *B. subtilis*. Top: *L. monocytogenes (Lm*) EGD-e (single chromosomal locus). Middle: *B. thuringiensis (Bt*) 407 (two chromosomal loci). Bottom: *B. subtilis (Bs*) 168. The *Bs* 168 gene annotations are in accordance with gene or protein names for the corresponding RefSeq entries. Where orthologs to the *Bs* 168 genes are present in *Bt* 407 and/or *Lm* EGD-e, as determined by reciprocal best BLAST hits, these are given the same gene names, except for *fliS, fliE, fliH,* og *hag/flaA/flaB* (see **Supplementary Table 1**). Annotation for the following genes lacking *Bs* 168 orthologs is according to the specified references: *mogR* (Grundling et al., 2004), *gmaR* (Shen et al., 2006), *flgE* (NP_464224 and ZP_04138734), and *fliN* (ZP_04138744). Hypothetical genes in *Lm* EGD-e and *Bt* 407 are indicated by locus tags. Protein orthologs or blocks of colinear orthologs are indicated by identical coloring in two or three strains. Genes shown in white do not have orthologs in any of the two other strains.

### 3.2 Non-motile *B. anthracis* and *B. pseudomycoides* have retained mogR

As has previously been noted (Klee et al., 2010), comparative alignments between the motility loci of different *B. cereus* group strains indicate that the flagellin gene locus is an evolutionary hotspot, as the number of flagellin genes in each strain varies between one and five copies, and several strains also contain additional sets of genes in this locus. Isolates belonging to the *B. cereus* group may be divided into seven phylogenetic subgroups (I to VII) and for which, with the exception of *B. pseudomycoides* (group I) and *B. cytotoxicus* (group VII), species designations do not strictly follow the phylogenetic group designations. Instead, isolates belonging to the different phylogenetic subgroups have distinct ecotypes and preferred growth temperature ranges (Guinebretière et al., 2008). An analysis of *B. cereus* group genomes revealed striking differences between the motility gene clusters in strains belonging to the various phylogenetic groups (**Figure 2**). The motility gene clusters of strains in phylogenetic subgroups III, IV and V were overall highly similar to that of *B. thuringiensis* 407, with a few exceptions mainly related to insertion of transposable elements and variation in the number of flagellin genes. Of note, strains of *B. anthracis,* which constitutes a highly clonal lineage within subgroup III, all carry an 11.5 kb insertion in place of the second flagellin gene (*flaB),* as well as nonsense mutations in a series of the conserved motility genes (*motP, cheA, flgL, fliF,* BAS1560 [ortholog to BTB_RS08355], *fliK,* BAS1570 [ortholog to BTB_RS08380], *cheV,* BAS1584 [ortholog to BTB_RS08410], *fliM,* and *flhF). B. anthracis* strains are thus non-motile, however still carry an intact *mogR* gene (**Figure 2**).

**Figure 2.**
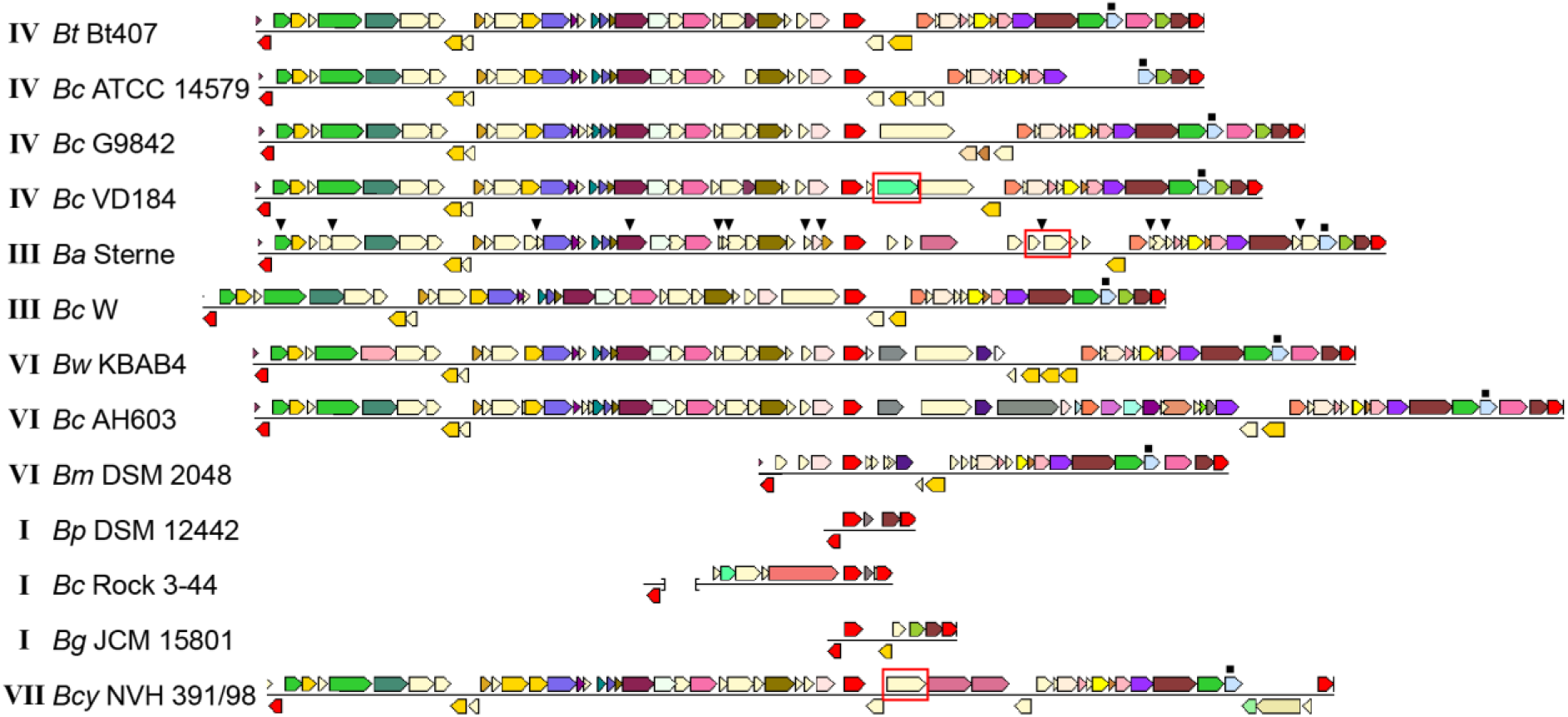
Comparison of motility gene loci in selected *Bacillus cereus* group strains. Loci from different strains are aligned with respect to *mogR,* shown in red in the central part of the alignment. For each strain, the first and last gene (shown in red) are conserved genes outside of the motility gene locus itself (orthologs to BTB_RS08235 and BTB_RS08480, respectively). These are included to indicate that the flanking regions and location of the motility locus are generally conserved between strains. The last gene of the intact locus with a predicted function mapping within motility, *flgG* (light blue), is indicated by a filled square above the gene. For genes shown in a color other than red, identical color indicates that genes are from the same orthologous group (top COG hit), except light yellow which indicates no COG assignment. The *gmaR* gene is indicated with a red box. In the *B. anthracis* strain, the filled triangles above genes indicate the location of nonsense or frameshift mutations. The brackets in the representation of the *Bc* Rock 3-44 sequence indicates that this locus is split between two contigs. Roman numerals next to the strain names indicate the phylogenetic subgroup within the *B. cereus* group to which the strain belongs (Guinebretière et al., 2008). *Bm* Rock 1-4, *Bm* Rock 3-17, *Bm* 219298 og *Bp* DSM 12442 from subgroup I had indentical loci; only *Bp* DSM 12442 is shown. The figure was assembled from images obtained from the Conserved Neighbourhood Viewer of the Integrated Microbial Genomes system (Markowitz et al., 2010), using *B. thuringiensis* 407 proteins BTB_RS08235, BTB_RS08390 (*mogR),* and BTB_RS08480 as query sequences. Abbreviations: *Bc*, *B. cereus; Bt, B. thuringiensis; Ba, B. anthracis; Bw, B. weihenstephanensis; Bm, B. mycoides; Bp, B. pseudomycoides; Bcy, B. cytotoxicus; Bg, Bacillus gaemokensis*.

Phylogenetic subgroup I corresponds to the species *B. pseudomycoides*, but also includes strains for which an original identification as *B. cereus* or *B. mycoides* has not yet been revised (Guinebretière et al., 2008). BLASTP searches were performed against available subgroup I proteomes using the corresponding protein sequences for the genes from the motility locus of *B. thuringiensis* 407 as query (**Figure 2**). Among six fully sequenced strains (single scaffold per replicon or closed genome) belonging to subgroup I (219298, BTZ, Rock 3-44, DSM 12442, Rock 1-4 and Rock 3-17), all contained a severely reduced motility gene cluster, and strikingly, the only conserved motility-related gene in the motility locus of these strains is *mogR* (**Figure 2**). The absence of all motility genes except *mogR* in all six fully sequenced group I strains analyzed, as well as in all except for one (AFS092012) of the 100 additional group I strains annotated as *B. pseudomycoides* at the NCBI and for which a genome sequence at scaffold level was available, indicates that absence of motility is likely a general characteristic of this genetic group, in line with the description of *B. pseudomycoides* as a nonmotile species (Nakamura, 1998). The predicted MogR proteins of the *B. pseudomycoides* group show ~56% and ~30% pairwise identity to the corresponding orthologs in *B. thuringiensis* 407 and *L. monocytogenes* EGD-e, respectively. A multiple sequence alignment demonstrated that the *L. monocytogenes* MogR residues predicted to make base-specific DNA contacts in *L. monocytogenes* (Shen et al., 2009) have been conserved in the MogR proteins from all examined *B. cereus* group strains, including those belonging to non-motile species (**Supplementary Figure 1**). Interestingly, an ortholog of *gmaR,* encoding a temperature-controlled antirepressor of MogR in *L. monocytogenes* (Kamp and Higgins, 2009), is found in *B. cytotoxicus* strains (cluster VII), but is absent in most *B. cereus* group strains belonging to other phylogenetic clusters.

### 3.3 Flagellin gene expression drops following a sharp increase in *mogR* expression

*B. thuringiensis* 407 produces two flagellins, structural proteins building the flagellum main filament, from chromosomal genes *flaA* and *flaB* (Lövgren et al., 1993). Expression of *mogR* (BTB_RS08390), *flaA* (BTB_RS08400) and *flaB* (BTB_RS08395) was followed at the transcriptional level throughout the bacterial growth phase using RT-qPCR. In contrast to previous proteomics studies which indicated that in *B. thuringiensis* 407 only the first flagellin gene (*flaA*) was expressed in early stationary phase (Gohar et al., 2005), results showed that both *flaA* and *flaB* are indeed expressed at the RNA level, although expression of *flaA* was approximately four-fold higher than that of *flaB* (**Figure 3**). The two *fla* genes reached maximum expression at 3.5 h, at the transition between exponential and stationary growth phase, in agreement with corresponding data obtained using a transcriptional fusion between *lacZ* and the *flaA* promoter (Houry et al., 2010). These data also corresponded well with microscopic observations of cultures sampled throughout the growth phase. The expression of *mogR* increased 64-fold between the 2.5 h and 3.5 h time-point (**Figure 3**), peaking at 4 h, at which time *flaA* and *flaB* expression was rapidly decreasing. This is in parallel with the expression pattern found for *mogR* in *B. anthracis* (Bergman et al., 2006). While downregulation of motility-related genes upon entry to the stationary phase is common in both Gram-negative and Gram-positive bacteria (Ramirez Santos et al., 2005; Han et al., 2017), it was nevertheless interesting to note that expression of the *flaA* and *flaB* genes dropped after 3.5 h, subsequent to the sharp increase in *mogR* transcription (**Figure 3**), in line with a role for MogR in repression of motility in *B. thuringiensis*.

**Figure 3.**
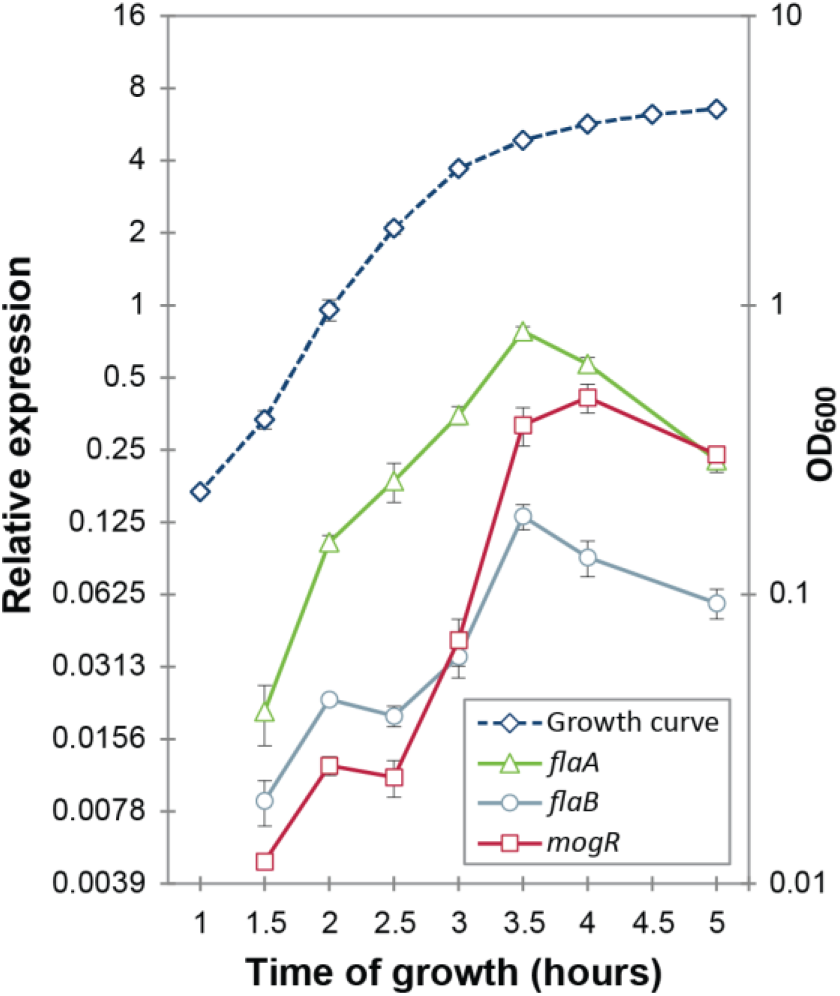
Gene expression analysis of *mogR* and flagellin genes *flaA* and *flaB* by RT-qPCR. The expression of each gene at each time-point was normalized to the expression of the reference genes *gatB*, *rpsU* and *udp*. Cultures were grown at 30°C in bactopeptone culture medium, with the growth curve plotted on the right-hand vertical axis. Averages and standard errors of the means from three independent experiments are shown.

### 3.4 Overexpression of MogR renders *B. thuringiensis* 407 non-flagellated and non-motile

A MogR overexpression strain was constructed using the pHT304-P_*xyl*_ expression vector carrying a xylose-inducible promoter, as well as an isogenic strain overexpressing a version of MogR where two mutations were introduced: Q119A and N120A (plasmids and strains used are listed in **Table 1**). In the mutated protein (referred to as MogR^QN→AA^), alanine substitutions were made for two of the six amino acids corresponding to residues in *L. monocytogenes* MogR that were shown to make base-specific contacts with DNA, in cognate DNA recognition sites positioned upstream of *L. monocytogenes* genes that were subject to direct MogR transcriptional repression (Shen et al., 2009; **Supplementary Figure 1**). Cellular growth rates were not affected by MogR or MogR^QN→AA^ overexpression (relative to an empty vector control) at 25°C, 30°C or 37°C (**Supplementary Figure 2**).

Motility of the MogR overexpression strain was investigated by swimming assays employing 0.3% LB agar plates, showing that cells overexpressing MogR were non-motile, and that the repression of motility was dependent on the two amino acid residues (Q119 and N120) mutated in MogR^QN→AA^, at 30°C (**Figures 4A,B**), as well as at 25°C and 37°C. Swimming motility was also monitored by light microscopy and found to be completely lost upon MogR overexpression at all time points and temperatures examined (every half hour between 1 and 7 h of cultivation, at 25°C, 30°C and 37°C). To determine whether the reduced motility was due to a loss of flagellar structure or flagellar rotation, strains were analyzed by Atomic Force Microscopy (AFM). AFM analyses showed that the MogR overexpression strain was completely devoid of flagellar structures, while overexpression of MogR^QN→AA^ restored the empty vector control phenotype (**Figures 4D-F**). In accordance, a Western blot of the MogR overexpression strain using anti-flagellin antibodies showed that flagellin proteins were below the level of detection in whole cell extracts and severely attenuated in outer cell surface extracts compared to in the vector control strain, while being expressed at comparable levels in the *B. thuringiensis* 407 wild type strain and in *B. thuringiensis* 407 overexpressing MogR^QN→AA^ (**Figure 4G**). In a Δ*flaAB* negative control strain in which the genes encoding the FlaA and FlaB flagellins had been deleted, no flagellin proteins could be detected neither in the whole cell extract nor on the cell surface. In *L. monocytogenes*, flagellin genes were previously found to be directly repressed by MogR, and in order to investigate if MogR function was conserved in *B. thuringiensis* 407, the expression of flagellin genes was investigated in the MogR overexpression strain relative to the empty vector and MogR^QN→AA^ control strains, by quantitative RT-PCR. A 30-fold (*flaA*) and 50-fold (*flaB*) reduction in transcription, respectively, was seen in the MogR overexpression strain relative to the empty vector control, while no difference was observed relative to the empty vector strain when overexpressing MogR^QN→AA^ (**Figure 4C**).

**Figure 4:**
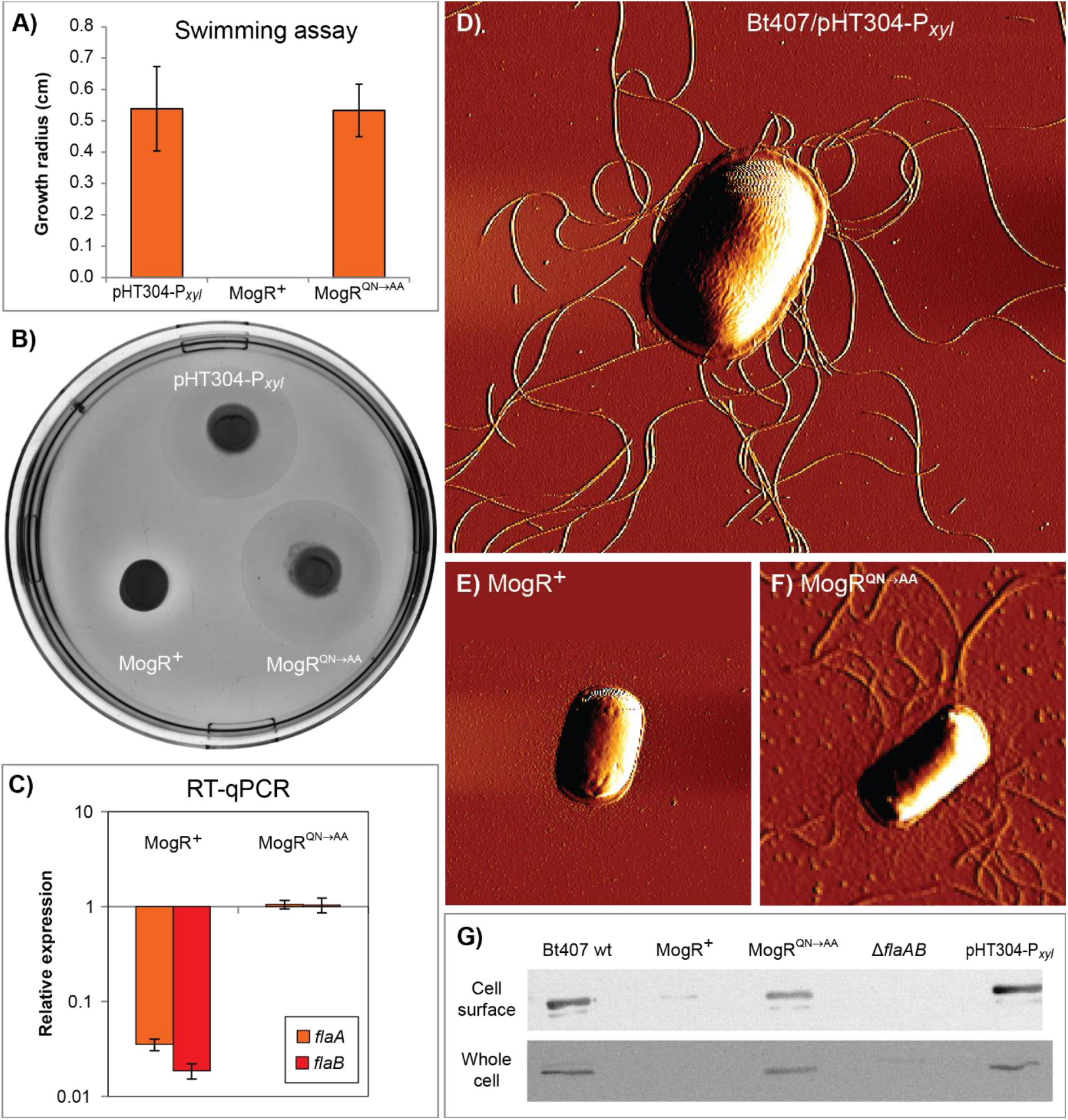
Analysis of motility and expression of flagella upon MogR overexpression in *B. thuringiensis* 407. Assays were conducted using the following strains: *B. thuringiensis* 407 pHT304-P_*xyl*_ empty vector control strain (pHT304-P*_xyl_*); *B. thuringiensis* 407 pHT304-P*_xyl_*-*mogR* (MogR^+^), overexpressing MogR; and *B. thuringiensis* 407 *pHT304-P_xyl_-mogR^QN→AA^* (MogR^QN→AA^), overexpressing a mutated form of MogR (see **Table 1**). Expression was induced with 1 mM xylose and cells were grown at 30°C. **(A,B)** Swimming motility was determined following growth on 0.3% LB agar for 7 h. In **(A)**, the mean and corresponding standard error values obtained from three independent experiments are shown. **(C)** Gene expression analysis of *flaA* and *flaB* by RT-qPCR. The relative expression of *flaA* and *flaB* in the MogR or MogR^QN→AA^ strains was normalized to the expression level of each respective gene in the empty vector control strain. Averages and standard errors of the means from three experiments are shown. **(D,E,F)** Atomic Force Microscopy images of bacterial cells grown to an OD_600_ of 3. **(G)** Western immunoblots showing the level of flagellin protein present in cell surface and whole cell extracts. Here, the *B. thuringiensis* 407 wild type (wt) and *B. thuringiensis* 407 Δ*flaAB* strains were included as controls.

### 3.5 Overexpression of MogR affects the expression of genes not related to motility functions

Global microarray-based transcriptional profiling was used to identify genes whose expression was affected by MogR overexpression in *B. thuringiensis* 407 relative to the empty vector control strain. Samples were taken in the early stationary growth phase (3 h), and a total of 110 genes were identified to be differentially regulated by MogR (FDR-corrected *p*<0.05), either directly or indirectly. Selected genes are listed in **Table 2** (see **Supplementary Table 3** for complete list). Of the differentially expressed genes, 87 were repressed by MogR overexpression, while 23 genes exhibited higher transcriptional levels compared to the control. In total, 21 motility genes were downregulated by MogR overexpression, including flagellin genes *flaA* and *flaB* (**Supplementary Figure 3**). Overexpression of MogR affected genes widely dispersed in the motility loci, including the *motAB* operon located separately in the chromosome. Interestingly, the microarray analysis revealed that MogR also affected the expression of 89 non-motility-related genes, including six virulence genes which were downregulated. The latter genes included *inhA* encoding immune inhibitor A, *hblA* and *hblD* encoding enterotoxin binding component B and lytic component L1, respectively, *sfp* encoding a subtilase family serine protease, as well as genes encoding phosphatidyl-choline specific phospholipase C and sphingomyelinase. Also downregulated were five stress-related genes: the *groES, groEL* and *grpE* chaperonin genes, *sigB* encoding the σ^B^ alternative sigma factor, and *hrcA*. The gene encoding the pleiotropic transcriptional regulator NprR (Dubois et al., 2012) was also downregulated. Among the 23 genes upregulated in the MogR overexpression strain were *sin*I, encoding a protein inducing biofilm formation by direct interaction and sequestration of the biofilm repressor SinR, and a putative capsular polysaccharide biosynthesis protein-encoding gene (BTB_RS26935) corresponding to BC5278 in *B. cereus* strain ATCC 14579. The genes BC5267 to BC5278 constitute a conserved locus in *B. cereus* and *B. thuringiensis* and are homologous to the *epsAO* locus of *B. subtilis* involved in the synthesis of the exopolysaccharide component of the biofilm matrix (Branda et al., 2001; Ivanova et al., 2003; Kearns et al., 2005; Fagerlund et al., 2014). Genes in this locus have been found to be implicated in biofilm formation in *B. cereus* in a pellicle biofilm model (Hayrapetyan et al., 2015; Okshevsky et al., 2017). Another upregulated gene was *cbpA*, encoding a collagen adhesion protein positively regulated by c-di-GMP (Finke et al., 2019).

**Table 2.** Genome-wide gene expression analysis of *B. thuringiensis* 407 overexpressing MogR relative to an empty vector control strain using oligonucleotide microarrays

| Gene | Locus tag in *B. thuringiensis* 407 | Locus tag in *B. cereus* ATCC 14579 | Predicted function | log<sub>2</sub>(fold change) in MogR overexpression strain relative to control |
| --- | --- | --- | --- | --- |
| **Motility-related gene function** |  |  |  |  |
| *cheA* | BTB_RS08255 | BC1628 | Chemotaxis protein, histidine kinase CheA | -1.80 |
| *flgK* | BTB_RS08295 | BC1636 | Flagellar hook-associated protein 1, FlgK | -1.67 |
| *fliS* | BTB_RS08310 | BC1639 | Flagellar biosynthesis protein FliS | -0.68 |
| *flgC* | BTB_RS08325 | BC1642 | Flagellar basal-body rod protein FlgC | -2.87 |
| *fliF* | BTB_RS08335 | BC1644 | Flagellar M-ring protein FliF | -2.31 |
| *fliG* | BTB_RS08340 | BC1645 | Flagellar motor switch protein FliG | -1.09 |
| *fliK* | BTB_RS08355 | BC1649 | Flagellar hook-length control protein FliK | -1.47 |
| *cheV* | BTB_RS08385 | BC1654 | Chemotaxis signal transduction protein CheV | -1.80 |
| *flaB* | BTB_RS08395 | BC1656 | Flagellin FlaB | -1.04 |
| *flaA* | BTB_RS08400 | BC1657 | Flagellin FlaA | -3.45 |
| *fliN* | BTB_RS08425 | BC1661 | Flagellar motor switch protein (fliN-homolog) | -0.99 |
| *fliP* | BTB_RS08430 | BC1665 | Flagellar biosynthesis protein FliP | -1.86 |
| *fliQ* | BTB_RS08435 | BC1666 | Flagellar export apparatus protein FliQ | -1.23 |
| *fliR* | BTB_RS08440 | BC1667 | Flagellar biosynthesis protein FliR | -1.02 |
| *flhB* | BTB_RS08445 | BC1668 | Flagellar biosynthesis protein FlhB | -1.60 |
| *flhA* | BTB_RS08450 | BC1669 | Flagellar biosynthesis protein FlhA | -1.43 |
| *motB* | BTB_RS22905 | BC4512 | Flagellar motor protein MotB (H<sup>+</sup>-coupled stator) | -3.01 |
| *motA* | BTB_RS22910 | BC4513 | Flagellar motor protein MotA (H<sup>+</sup>-coupled stator) | -1.74 |
| *mcpA* | BTB_RS02175 | BC0422 | Methyl-accepting chemotaxis protein (carrying upstream "off" c-di-GMP riboswitch) | -1.31 |
| **Virulence-related gene function** |  |  |  |  |
| *nprR* | BTB_RS03040 | BC0598 | Transcriptional activator NprR (necrotrophism regulator) | -0.89 |
| *inhA* | BTB_RS03410 | BC0666 | Immune inhibitor A | -1.03 |
| *plcB* | BTB_RS03430 | BC0670 | Phospholipase C (PC-PLC) | -2.32 |
| *sph* | BTB_RS03435 | BC0671 | Sphingomyelinase C (Smase) | -1.47 |
| *hblA* | BTB_RS12545 | BC3102 | Hbl component B | -1.13 |
| *hblD* | BTB_RS12540 | BC3103 | Hbl component L1 | -1.59 |
| *sfp* | BTB_RS18825 | BC3762 | Serine protease (subtilase family) | -0.84 |
| *cbpA* | BTB_RS05575 | BC1060 | Collagen adhesion protein (carrying upstream c-di-GMP "on" riboswitch) | 3.02 |
| **Putative biofilm-related gene function** |  |  |  |  |
| *sinI* | BTB_RS06545 | BC1283 | SinI biofilm repressor antagonist | 1.08 |
| *ywgC* | BTB_RS26935 | BC5278 | Putative capsular polysaccharide biosynthesis protein | 0.71 |

### 3.6 Q119 and N120 are critical for MogR binding to specific sites in the *B. thuringiensis* 407 genome *in vitro*

DNA regions upstream of genes differentially expressed in the *B. thuringiensis* 407 MogR overexpression strain were investigated for candidate MogR binding sites, based on sequence searches using the *L. monocytogenes* consensus MogR binding sequence (TTTTWWNWWAAAA; Shen et al., 2009). Three putative binding sites were predicted in the intergenic region upstream of the flagellin (*flaAB*) operon, two potential sites upstream of *cheY,* while four putative binding sites were predicted in the promoter region of the *hbl* operon (Lindbäck et al., 1999), all overlapping the −10 promoter element (**Figure 5A**). Among the other genes downregulated by MogR overexpression, multiple binding sites were found upstream of *inhA,* as well as upstream of the biofilm-promoting gene *sin*I and *mogR* itself.

**Figure 5:**
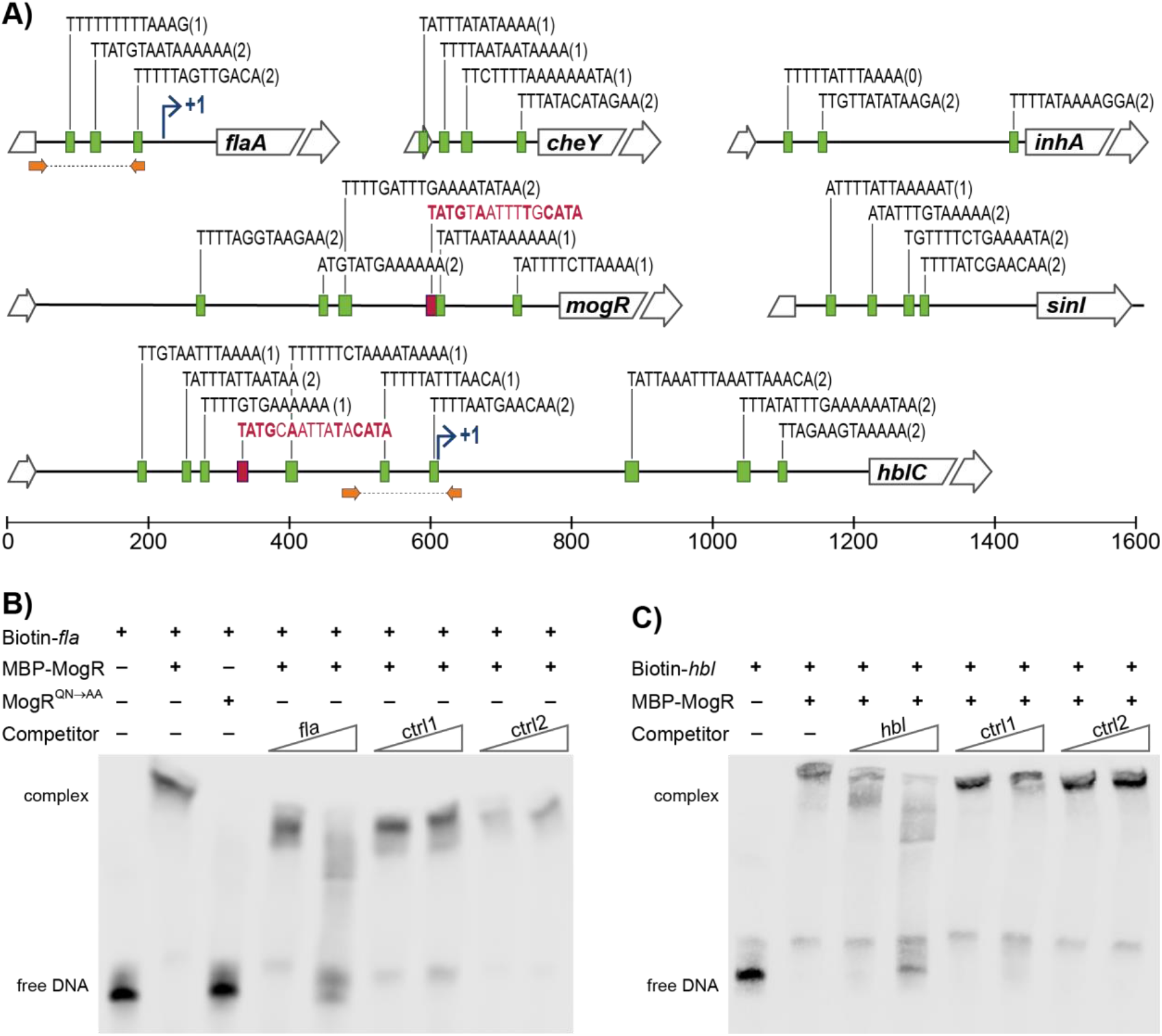
MogR binding to promoter regions in *B. thuringiensis* 407. **(A)** Putative MogR binding sites (indicated by green boxes) in the intergenic regions upstream of the *flaA, cheY, inhA, mogR, sin*I and *hblC* genes, allowing up to two mismatches from the sequence TTTTWWNWWAAAA, with the number of mismatches in parentheses. Putative PlcR boxes, TATGNANNNNTNCATA, are included in red (only perfect matches included). Broken arrows indicates transcriptional start sites where applicable. Location of primers used for generating *fla* and *hbl* promoter sequence probes for the electrophoretic mobility shift assay (EMSA) in **(B,C)** are indicated with orange arrows. **(B,C)** EMSAs of purified MBP-MogR fusion protein binding to purified PCR products representing the promoter regions of *flaA* **(B)** and *hblC* **(C)**, as indicated in **(A)**. Biotin-labeled *flaA* or *hblC* DNA fragments were incubated with 1 μg (equivalent to 1.4 μM) purified MBP-MogR protein, in the presence or absence of unlabeled competitor DNA at 50-fold or 1000-fold molar excess: *fla* or *hbl* fragment, and nonspecific competitors ctrl1 (*B. thuringiensis* 407 BTB_RS08990) and ctrl2 (gene encoding Epstein Barr virus nuclear antigen). In **(B)**, 1μg MBP-MogR^QN→AA^ was used for DNA binding, as a control.

To determine whether *B. thuringiensis* MogR can function as a DNA-binding protein, and investigate whether MogR may affect the expression of candidate genes in *B. thuringiensis* 407 by directly binding to upstream regulatory regions, we carried out electrophoretic mobility shift assays (EMSA) with a purified maltose-binding protein (MBP)-MogR fusion protein, and purified DNA fragments constituting PCR-amplified candidate regulatory regions (**Figure 5A**). Two promoter regions upstream of genes shown to be differentially expressed in the microarray experiments were selected for analysis. Results showed that MogR could bind the promoter regions upstream of both *flaAB* and the *hbl* enterotoxin locus *in vitro* (**Figures 5B,C**). The amount of MogR protein to be used in the assay had been optimized by titration in a prior experiment (**Supplementary Figure 4**). Importantly, DNA binding was abolished when the experiment was performed using MogR^QN→AA^, in which the conserved Q119 and N120 residues shown to make base-specific contacts with the MogR recognition site in *L. monocytogenes* (Shen et al., 2009) had been substituted (**Figure 5B**). The binding reactions to which competing unlabelled DNA were added (labelled ctrl1 and ctrl2 in **Figures 5B,C**) indicated that MogR exhibits sequence specific binding to *flaAB* and *hbl* upstream regions *in vitro*.

### 3.7 Cytotoxin production and virulence are reduced upon MogR overexpression

Western blot analyses were performed to measure the effect of MogR overexpression on the amount of secreted enterotoxins. The genes encoding the Hbl, CytK and Nhe enterotoxins are all part of the PlcR regulon (Gohar et al., 2008), and thus commonly co-regulated in *B. cereus* group bacteria. Therefore, the expression of all three was examined, despite the fact that an upstream promoter containing putative MogR binding sites and differential expression upon MogR overexpression was only observed for the genes encoding Hbl. All three enterotoxins were present but at clearly reduced levels in the culture supernatant from the MogR overexpression strain, compared to that of empty vector and MogR^QN→AA^ control strains (**Figure 6A**). To further assess the effect of MogR on virulence capacity, supernatants collected from mutant and overexpression strains grown to exponential phase were tested in an *in vitro* Vero cell toxicity assay (Lindbäck and Granum, 2006). Supernatants from the MogR overexpressing strain, which synthesized lower amounts of the Hbl, Nhe and CytK toxins (**Figure 6A**), were less cytotoxic than those of the empty vector control (*p*=0,059) and the MogR^QN→AA^ strain (**Figure 6B**). In order to more confidently assess the effect of MogR on virulence, an *in vivo* model was used, where the strains were tested in an oral infection model using *Galleria mellonella* larvae. In accordance with the *in vitro* toxicity data, the MogR overexpressing strain was severely attenuated in virulence compared to the empty vector control strain *in vivo* (*p*=0.02; **Figure 6C**).

**Figure 6.**
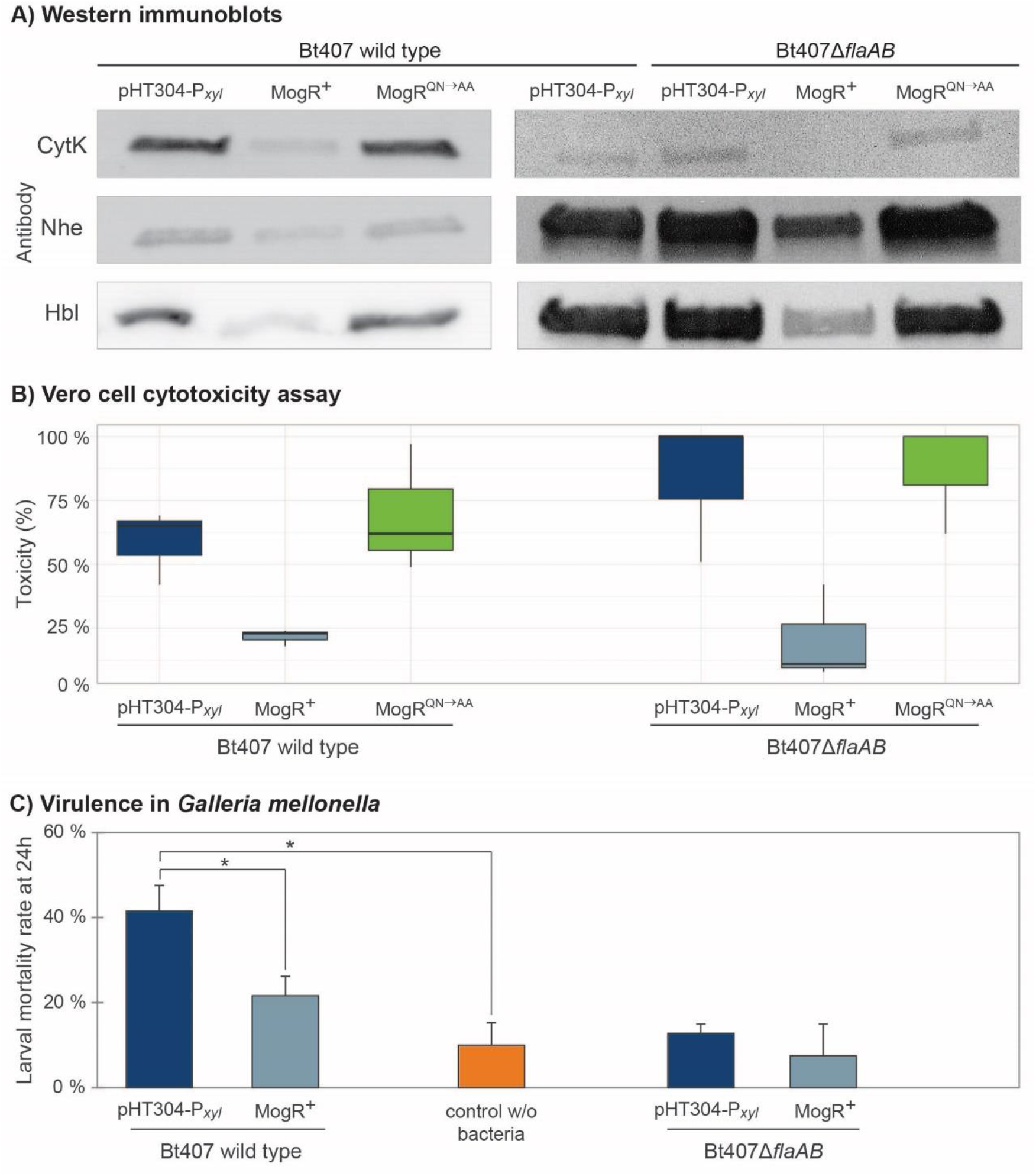
Analysis of the effect of overexpression of MogR on virulence and toxicity phenotypes. **(A)** Immunoblots against CytK, Nhe and Hbl enterotoxins using culture supernatants from *B. thuringiensis* 407 carrying the empty vector (pHT304-P*_xyl_*), a MogR overexpression strain (MogR^+^), and a strain overexpressing a mutated form of MogR (MogR^QN→AA^). Constructs were analysed in *B. thuringiensis* 407 wild type and *flaAB* deletion (Δ*flaAB*) backgrounds. Cultures were grown for 4.5 h with 1 mM xylose induction prior to harvest. The experiments were repeated at least three times and representative blots are shown. **(B)** Vero cell cytotoxicity assay using supernatants from cultures of the same strains as in **(A)**. The median and first and third quartiles (25^th^ percentile and 75^th^ percentile) of three independent experiments are shown, as well as whiskers indicating the data variability outside the upper and lower quartiles. **(C)** Virulencein *Galleria mellonella* insect larvae (oral force-feeding infection model), using empty vector and MogR overexpression strains, in *B. thuringiensis* 407 wild type and Δ*flaAB* backgrounds, respectively. Bacteria were in all cases mixed with Cry 1C toxin and xylose. Also, a control with no bacteria but fed with Cry toxin and xylose was included. The means and standard errors of the mean (bars) of seven independent experiments are shown. Statistically significant differences (Tukey’s *post hoc* test, *p*<0.02) are indicated with asterisks (*).

### 3.8 Biofilm formation is increased upon MogR overexpression in a *flaAB* negative background

The MogR overexpression strain was compared to the empty vector and MogR^QN→AA^ strains both in glass tube and microtiter plate biofilm assays (**Figures 7A,B**). The glass tube assay showed similar development of a biofilm pellicle at the air-liquid surface in all three strains: After 72 h the pellicle in the MogR overexpression strain was only slightly (14%) thicker than that of the empty vector and MogR^QN→AA^ strains (**Figure 7A**). In the microtiter plate assay, no significant effect of MogR overexpression on biofilm formation was observed at 48 or 72 h, with a slight decrease in biofilm formation at 24 h compared to MogR^QN→AA^ and empty vector controls (**Figure 7B**). Overexpression in a Δ*flaAB* background however allowed examination of the effect of the MogR protein on biofilm formation, independently of any effect conferred through flagella. Interestingly, in the Δ*flaAB* host, overexpression of MogR led to a substantial increase in biofilm formation relative to the MogR^QN→AA^ and empty vector controls (**Figure 7C**), while biofilm formation in the control strain overexpressing MogR^QN→AA^ was comparable to that of the strain carrying empty vector. Intriguingly, LC-MS/MS quantitation of the biofilm-related second messenger cyclic di-GMP (c-di-GMP) in cellular extracts indicated slightly increased levels of c-di-GMP in the MogR overexpression strain (wild type background; three measurements: 8.38 ng mL^−1^; 6.0 ng mL^−1^; 9.0 ng mL^−1^) compared to the *B. thuringiensis* 407 empty vector strain (two measurements: < LOD; 6.0 ng mL^−1^) (Limit of detection, LOD: 0.8 ng mL^−1^; Limit of quantitation, LOQ: 3.5 ng mL^−1^).

**Figure 7.**
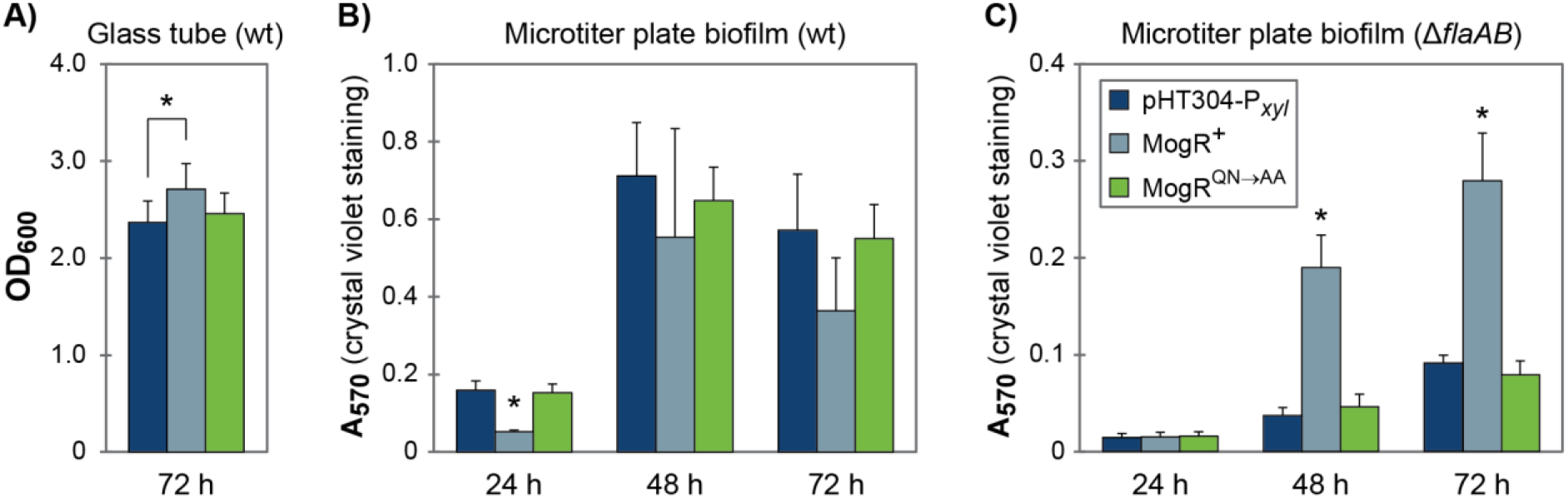
MogR overexpression affects biofilm formation. **(A)** Biofilm formation in glass tubes after 72 h of growth. **(B,C)** Biofilm formation in a crystal violet microtiter plate assay after 24, 48, and 72 h of growth. The legend shown in **(C)** also applies to figures **(A,B)**. The means and standard error of the means for **(A)** five or **(B, C)** three independent experiments is shown. Statistically significant differences (within each time point; Tukey’s *post hoc* test, *p*<0.05) are indicated with an asterisk (*); in **(B,C)** the asterisk (*) indicates that the MogR^+^ strain demonstrates a statistically significant difference to both the empty vector control strain and the overexpression of a mutated form of MogR (MogR^QN→AA^). In **(A)**, only the MogR^+^ and empty vector control strains (pHT304-P_*xyl*_) were significantly different.

## 4 Discussion

Overexpression of MogR leads to repression of a range of motility genes in *B. thuringiensis* 407, including the flagellins, rendering the bacteria non-flagellated and non-motile. In addition, amino acids in *B. thuringensis* MogR which correspond to those shown to mediate base-specific DNA contacts in *L. monocytogenes* MogR are clearly essential for *B. thuringiensis* MogR DNA binding *in vitro*. In accordance, overexpression of the mutated variant of MogR restores phenotypes to those of a vector control strain. Collectively these results, and analyses of MogR binding to predicted regulatory sites *in vitro,* strongly support the role of *B. thuringiensis* MogR as a DNA-binding transcriptional regulator and that the role of MogR as a key motility regulator in *B. thuringiensis* is conserved with that in *L. monocytogenes*. Global gene expression profiling however showed that in *B. thuringiensis* a large number of non-motility genes were also affected by MogR overexpression, including genes involved in stress response, virulence and biofilm formation. This was paralleled by an observed decrease in cytotoxin production, Vero cell toxicity and virulence in the *G. mellonella* infection model, indicating that MogR can affect additional phenotypes in *B. cereus* group bacteria. This is in apparent contrast to *L. monocytogenes,* where the primary function of MogR during intracellular infection is to repress flagellar motility gene expression, and where deletion of the flagellin gene (*flaA*) in a *mogR* deletion strain (producing Δ*mogR* Δ*flaA*) restores the LD_50_ values to wild type levels in a mouse model of infection (Shen and Higgins, 2006).

In bacteria, motility and virulence can be subject to coordinated downregulation, e.g. as is observed for intracellular signaling networks involving the second messenger c-di-GMP (Fagerlund et al., 2016). Flagellar motility and virulence appears to be co-regulated in *B. cereus* group bacteria (Zhang et al., 1993; Ghelardi et al., 2002; Bouillaut et al., 2005; Ghelardi et al., 2007a; Fagerlund et al., 2010; Salvetti et al., 2011; Mazzantini et al., 2016). For instance, a *B. cereus* strain deleted in *flhA,* which encodes a component of the flagellum-formation apparatus, is impaired in adhesion (Ramarao and Lereclus, 2006), expression of virulence factors (Ghelardi et al., 2002; Fagerlund et al., 2010), and in cytotoxicity and virulence independently of PlcR (Bouillaut et al., 2005), which is the principal transcriptional activator of *B. cereus* virulence gene expression. Similarly, deletion of the gene encoding the signal recognition particle-like GTPase FlhF, located directly downstream of *flhA,* resulted in a marked reduction in the number of flagella, and a reduction in secretion of several virulence-related proteins with attenuation of virulence (Salvetti et al., 2007; Mazzantini et al., 2016; Mazzantini et al., 2020). A link between synthesis of the flagellar apparatus and the production of virulence factors has also been observed for *flhA* deletion mutants in other bacteria that do not carry a *mogR* ortholog (Fleiszig et al., 2001; Carrillo et al., 2004). However, the effect on toxin production observed following deletion of *flhA* or *flhF* was clearly not replicated in the *flaAB* deletion mutant, showing that not all deletions in the motility locus affect toxin synthesis and/or transport.

The observed effects of MogR on toxin production and virulence-related phenotypes could be conferred through direct regulation, as potentially indicated through *in vitro* binding to the *hbl* upstream region, or indirectly through repression of genes in the motility locus, such as e.g. *flhA* and/or *flhF*, or both. As *plcR* expression was not affected by MogR overexpression, and only six out of the 28 genes comprising the PlcR regulon (Gohar et al., 2008) were differentially expressed in the microarray analysis, the observed repression of virulence by MogR is probably not due to direct repression of the virulence gene activator PlcR. Conversely, MogR was not identified as a member of the PlcR regulon, despite *in silico* analyses identifying a PlcR box upstream of *mogR* (Gohar et al., 2008; **Figure 5A**). However, it is possible that crosstalk between the MogR and PlcR regulators may nevertheless occur, at least under certain circumstances, as in many cases microarray experiments conducted under a given set of conditions fail to identify all members of a regulon. This was for example the case for the transcriptional regulator NprR, which was initially not considered part of the PlcR regulon despite the presence of a PlcR box in the promoter region (Gohar et al., 2008), but which was nevertheless later found to be positively controlled by PlcR (Dubois et al., 2013). In the current study, NprR was repressed by MogR overexpression. Like for virulence, motility also appears to be oppositely regulated by MogR and PlcR, as motility and motility gene expression was reduced in a *plcR* mutant (Gohar et al., 2002; Gohar et al., 2008). The existence and nature of putative molecular crosstalk mechanisms involving PlcR and MogR will require further study.

When employing a Δ*flaAB* genetic background to compare the effect of MogR overexpression on biofilm formation, relative to empty vector and MogR^QN→AA^ overexpression controls, a significant increase in biofilm formation was observed by MogR overexpression. In line with this, the transcriptional profiling experiments showed upregulated expression of the biofilm anti-repressor *sinI,* and BTB_RS26935 located in a putative polysaccharide locus previously shown to be implicated in the formation of the biofilm matrix in some *B. cereus* strains (Hayrapetyan et al., 2015; Okshevsky et al., 2017). Moreover, LC-MS/MS quantitation indicated a slight increase in cellular c-di-GMP upon MogR overexpression, which is linked to increased biofilm formation in *B. thuringiensis* 407 (Fagerlund et al., 2016; Fu et al., 2018). A microarray experiment analyzing the effect of *B. thuringiensis* 407 overexpressing the c-di-GMP synthesizing protein CdgF relative to an empty vector control strain (Fagerlund et al., 2016; A. Fagerlund, unpublished data deposited in the ArrayExpress database, accession no. E-MTAB-8898) showed that the expression of the biofilm regulators *spo0A* and *sin*I were upregulated and the transcriptional regulator *codY* was downregulated at high levels of c-di-GMP, all paralleling an increase in biofilm formation (Pflughoeft et al., 2011; Lindbäck et al., 2012; Fagerlund et al., 2014). Taken together, these experiments provide strong indication that MogR, directly or indirectly promotes biofilm formation in *B. thuringiensis* 407, a phenotype in line with the role of the protein in promoting sessility and repressing toxicity.

In *L. monocytogenes,* MogR activity is controlled by its anti-repressor GmaR (Kamp and Higgins, 2009). GmaR expression is halted when *L. monocytogenes* experiences 37°C, which allows MogR to repress flagellar synthesis during human infection. The *gmaR* gene is not present in most *B. cereus* group genomes, in line with *mogR* appearing to be regulated in a growth-dependent but temperature-independent manner in *B. thuringiensis* 407. It is however interesting to note that a *gmaR* ortholog is found in at least twelve sequenced *B. cereus* group strains, including all hitherto characterized strains of the thermotolerant species *B. cytotoxicus*, perhaps reflecting a need for strains in this phylogenetic group (group VII) to regulate flagellin synthesis at different temperatures. Most *B. anthracis* strains also harbor a copy of *gmaR,* which however is truncated and rendered non-functional due to mutation. Uncharacteristically, an apparently full-length *gmaR* copy was identified in the genome of *B. cereus* VD184, an environmental soil isolate belonging to phylogenetic cluster IV (Hoton et al., 2009; Jalasvuori et al., 2013), as well as *B. cereus* strains RU36C, F528-94, MB.22, B4118 and FSL W8-0932, and *Bacillus wiedmannii* FSL P4-0569, for which optimal growth temperatures are not known.

In conclusion, MogR is a key regulator of motility in *B. thuringiensis* 407, a function which is probably conserved across the motile species in the *B. cereus* group. Most importantly however, both microarray experiments, protein expression analyses, biofilm screening results, and the fact that the comparative genomics analyses showed *mogR* to be evolutionary retained in the non-motile species *B. mycoides*, *B. pseudomycoides*, and *B. anthracis*, as well as expressed in *B. anthracis*, speaks to the MogR transcriptional regulator serving additional functions in *B. cereus* group bacteria, beyond those identified in *L. monocytogenes*. In this respect it is interesting to note that all attempts to produce a gene deletion mutant in *mogR* in *B. thuringiensis* 407 were unsuccessful, and we cannot rule out the possibility of *mogR* constituting an essential gene in the *B. cereus* group. From the analyses performed in this study, it is tempting to speculate that MogR may serve as part of a molecular system coordinating motility, virulence, and biofilm formation, and possibly additional phenotypes. Biofilm formation and motility are reciprocally regulated in a number of bacterial species, including *B. cereus* group bacteria, by the second messenger c-di-GMP (Hengge, 2009; Fagerlund et al., 2016; Jenal et al., 2017; Fu et al., 2018), and several transcriptional regulators in both the *B. cereus* group and *L. monocytogenes* are known to have overlapping functions in regulation of virulence and biofilm formation (Hsueh et al., 2006; Lemon et al., 2010; Pflughoeft et al., 2011; Frenzel et al., 2012; Lindbäck et al., 2012; Fagerlund et al., 2014; Slamti et al., 2015; Böhm et al., 2016). It thus appears that, in the *B. cereus* group, there is substantial cross-regulation between motility, virulence, and biofilm formation, involving a multitude of regulatory networks based on both c-di-GMP and transcription factors (PlcR, MogR, NprR).

## 5 Data availability statement

The microarray data comparing overexpression of MogR with the empty vector control is available in the ArrayExpress database (www.ebi.ac.uk/arrayexpress) under accession number E-MTAB-7633, and the CdgF overexpression experiment has accession number E-MTAB-8898.

## Supporting information

Suppmentary Material File

## 6 Author Contributions

OØ and AF contributed to conception and design of the study. VS performed EMSA, motility assays, Western immunoblotting, and biofilm assays. VS and AF performed RT-qPCR. VS, MJ and AF performed cloning and mutagenesis. MJ performed the microarray experiment, SF performed growth experiments, IH performed AFM, TL performed cytotoxicity assays, and CN-L performed the *G. mellonella* infection experiments. AF performed bioinformatic, microarray, and statistical analyses and NT performed bioinformatic analysis of the subgroup I genomes. VS, OØ, and AF wrote the first draft of the manuscript. TL, IH, and CN-L wrote sections of the manuscript. All authors contributed to data analysis, read and approved the final manuscript.

## 9 Acknowledgements

We are very grateful to Erwin Märtlbauer, Ludwig-Maximilians-Universität, Germany, for monoclonal antibodies against Hbl and Nhe, and to Cecilie From, Norwegian University of Life Sciences, Norway, for anti-flagellin antibodies. We gratefully thank Didier Lereclus, INRA, France for the *E. coli/Bacillus* shuttle vector pHT304-P_*xyl*_, and Michel Gohar, INRA, France, for the *B. thuringiensis* 407 Δ*flaAB* strain. We thank Christophe Buisson, INRA, France, for performing force-feeding of insect larvae, and Marthe P. Parmer and Leon Reubsaet, School of Pharmacy, University of Oslo, Norway, for quantitation of cellular c-di-GMP levels.

