## Supplementary material for "MogR is a ubiquitous transcriptional repressor affecting motility, biofilm formation and virulence in *Bacillus thuringiensis*": Suppmentary Material File

### 1 Supplementary Figures

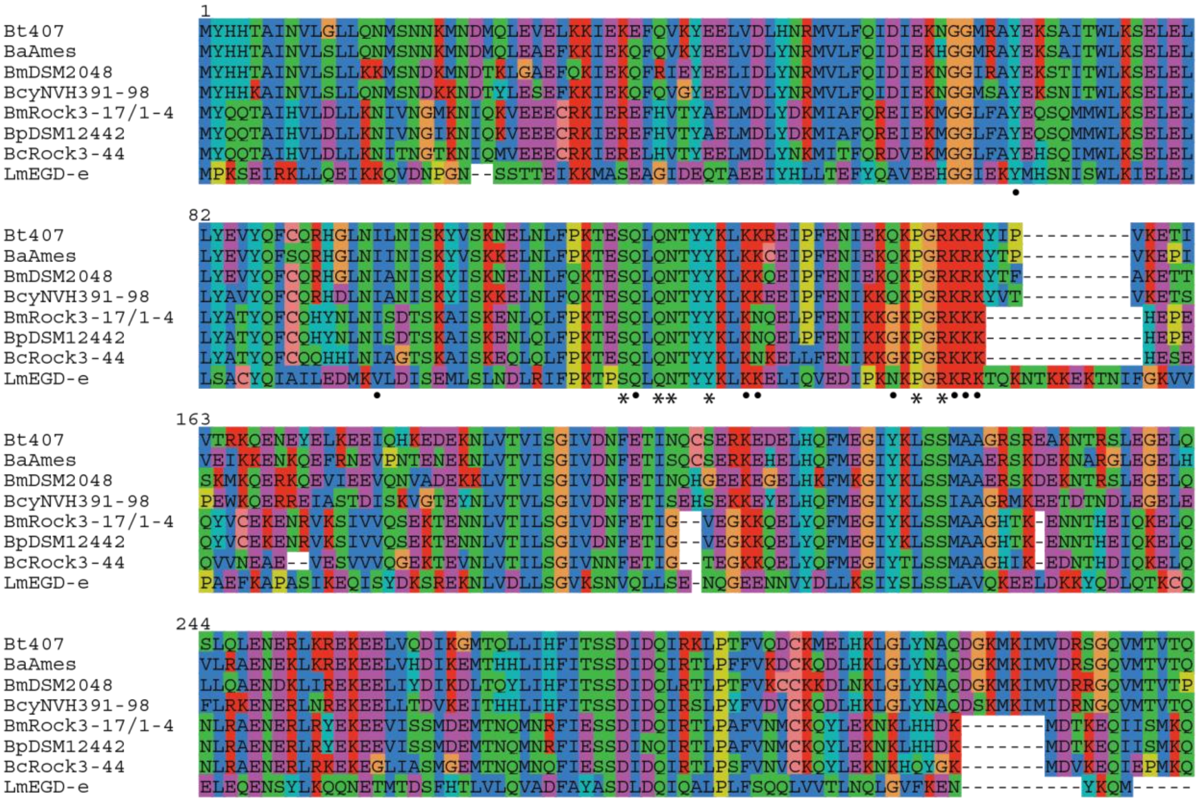


**Supplementary Figure 1. Multiple sequence alignment of MogR protein sequences.** MogR sequences were extracted from *L. monocytogenes* strain EGD-e (locus tag: lmo0674), *B. cytotoxicus* NVH 391/98 (Bcer98_1372), and *B. thuringiensis* 407 (BTB_RS08390), in addition to the following predicted non-motile strains: The group I strains *B. cereus* Rock 3-44 (bcere0022_13670), *B. pseudomycoides* DSM 12442 (bpmyx0001_15070) and *B. mycoides* Rock 3-17 and Rock 1-4 (bmyco0002_14720 and bmyco0003_14260, which are identical), the *B. anthracis* Ames strain (BA_1692), and the group VI strain *B. mycoides* DSM 2048 (bmyco0001_14970). The annotated start site of the MogR protein from *B. mycoides* DSM 2048 was adjusted to fit the start site predicted by EasyGene (Larsen and Krogh, 2003). The N-terminal amino acid sequence MLSGSEYK of MogR from the group VI *B. mycoides* strains and *B. pseudomycoides* DSM 12442 were trimmed from the alignment. Residues making DNA backbone contacts and those making base specific DNA contacts, as determined for *L. monocytogenes* MogR by Shen et al. (2009), are marked with a solid dot and a star, respectively.

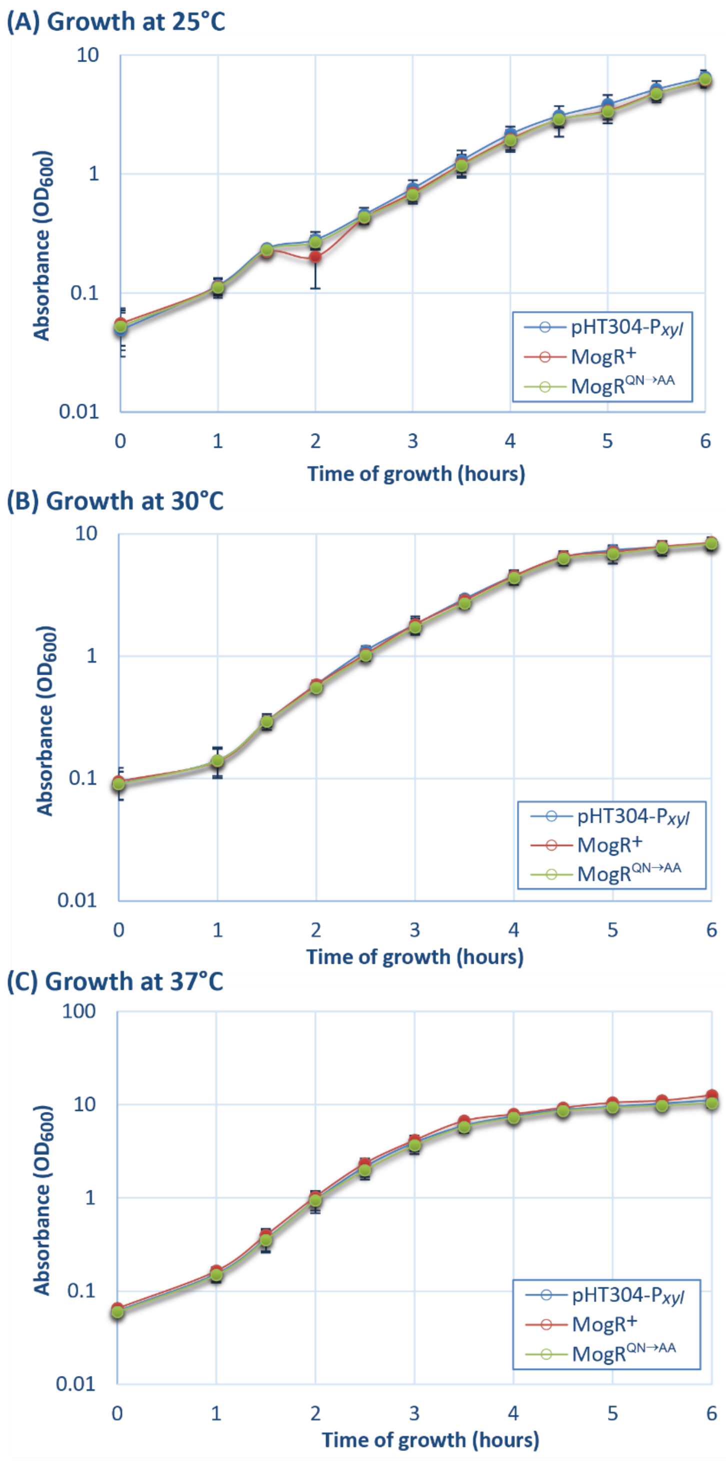


**Supplementary Figure 2. Growth curves at A) 25 °C, B) 30 °C and C) 37 °C.** Assays were conducted using *B. thuringiensis* 407 pHT304-$P_{xyl}$, an empty vector control strain (pHT304-$P_{xyl}$); *B. thuringiensis* 407 pHT304-$P_{xyl}$-*mogR* (MogR$^{+}$), a strain overexpressing MogR from the pHT304-$P_{xyl}$ plasmid vector; and *B. thuringiensis* 407 pHT304-$P_{xyl}$-*mogR*$^{QN \rightarrow AA}$ (MogR$^{QN \rightarrow AA}$), a strain overexpressing a mutated form of MogR from the pHT304-$P_{xyl}$ vector.

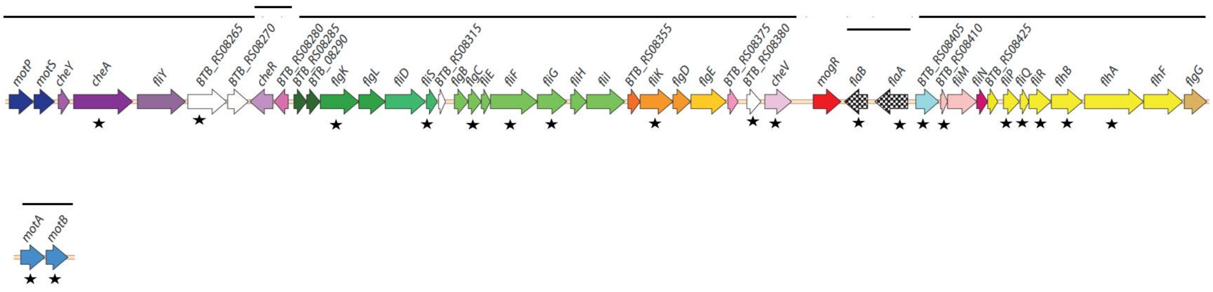


**Supplementary Figure 3. Graphic representation of genes in the motility loci from *B. thuringiensis* 407.** Solid lines above genes indicate operon structures, predicted from RNA-Seq data obtained from *B. cereus* strains ATCC 10987 and ATCC 14579 (Kristoffersen et al., 2012). Annotated genes are indicated with gene names positioned above the arrows. Genes affected by MogR overexpression are indicated by an asterisk.

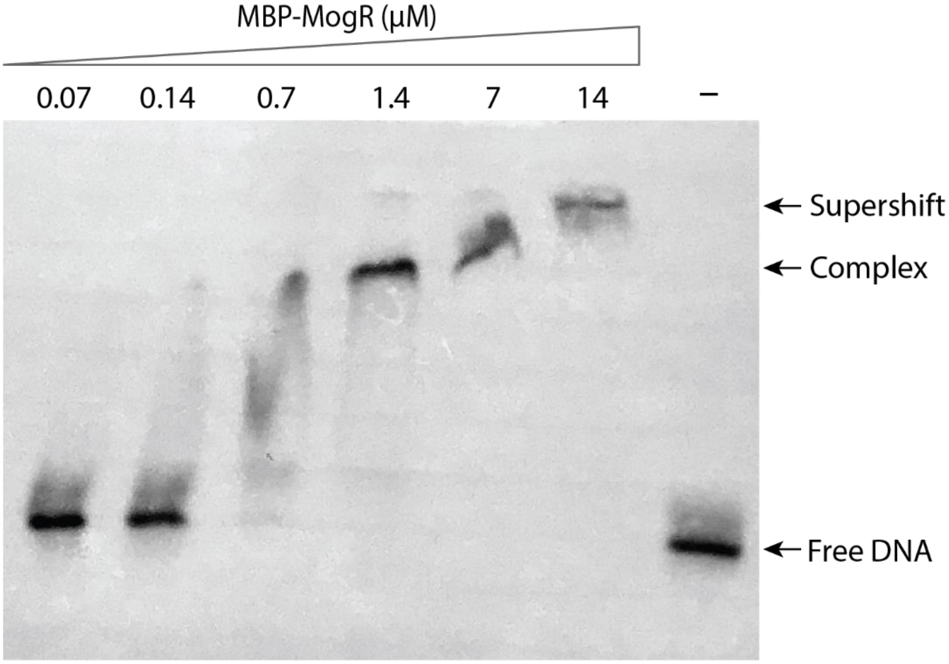


**Supplementary Figure 4. EMSA showing titration of MogR protein.** The experiment was carried out with biotin-labeled *fla* promoter region DNA and increasing concentrations of purified MBP-MogR fusion protein. A concentration of 1.4 µM is equivalent to 1 µg of the 37 kDa MBP-MogR protein in the binding reaction. The positions of free labelled DNA probe, shifted protein-DNA complex and supershifted protein-DNA complex are indicated. See Figure 5A for location of primers used for generating the *fla* promoter probe.

### 2 Supplementary Tables

**Supplementary Table 1. Reciprocal best BLAST hit analysis for motility proteins.**
Genomes of the following strains were subject to analyses: *B. thuringiensis* 407, *L. monocytogenes* EGD-e, and *B. subtilis* 168.

| | | Locus tags and seqIDs. Reference proteins are in shown in bold | | | Reciprocal best Blast hits* | | |
|---|---|---|---|---|---|---|---|
| Function / gene product | gene | Bt407 | EGDe | Bs168 | Bt407 vs EGDe | Bt407 vs Bs168 | EGDe vs Bs168 |
| Flagellar motor protein MotP ($Na^+$-coupled stator) | *motP* | **BTB_RS08240** | | BSU29730_ ytxD | no | yes (37%/92%) | no |
| Flagellar motor protein MotS ($Na^+$-coupled stator) | *motS* | **BTB_RS08245** | | BSU29720_ ytxE | no | yes (39%/95%) | no |
| Chemotaxis protein, response regulator CheY | *cheY* | **BTB_RS08250** | **lmo0691_cheY** | **BSU16330_cheY** | yes (75%/98%) | yes (68%/100%) | yes (69%/95%) |
| Chemotaxis protein, histidine kinase CheA | *cheA* | **BTB_RS08255** | **lmo0692_cheA** | **BSU16430_cheA** | yes (56%/105%) | yes (44%7103%) | yes (45%/103%) |
| Flagellar motor switch FliY | *fliY* | **BTB_RS08260** | **lmo0700** | **BSU16320_fliY** | yes (42%/99%) | yes (35%/76%) | yes (38%/65%) |
| hypothetical protein | | **BTB_RS08265** | | | no | no | |
| hypothetical protein | | **BTB_RS08270** | | | no | no | |
| Chemotaxis protein, methyltransferase CheR | *cheR* | **BTB_RS08275** | **lmo0683** | BSU22720_cheR | yes (44%/98%) | yes (48%/98%) | yes (47%/98%) |
| hypothetical protein | | **BTB_RS08280** | **lmo0687** | | yes (43%/100%) | no | no |
| Protein of unknown function (DUF327) | | **BTB_RS08285** | **lmo0703** | BSU00300_yaaR | yes (48%/93%) | yes (25%/89%) | yes (24%/55%) |
| FlgN-like superfamily protein | | **BTB_RS08290** | | | no | No | |
| Flagellar hook-associated protein FlgK | *flgK* | **BTB_RS08295_flgK** | **lmo0705_flgK** | **BSU35410_flgK** | yes (35%/62%) | yes (27%/47%) | yes (22%/106%) |
| Flagellar hook-associated protein FlgL | *flgL* | **BTB_RS08300_flgL** | **lmo0706_flgL** | **BSU35400_flgL** | yes (30%/101%) | yes (27%/93%) | yes (28%/69%) |
| Flagellar hook-associated protein FliD | *fliD* | **BTB_RS08305_fliD** | **lmo0707_fliD** | **BSU35340_fliD** | yes (28%/97%) | yes (24%/81%) | yes (24%/107%) |
| Flagellar protein FliS | *fliS* | **BTB_RS08310** | **lmo0708** | **BSU35330_fliS** | yes (54%/97%) | no | yes (21%/96%) |
| hypothetical protein | | **BTB_RS08315** | **lmo0709** | | no** | no | no |
| Flagellar basal-body rod protein FlgB | *flgB* | **BTB_RS08320_flgB** | **lmo0710_flgB** | **BSU16180_flgB** | yes (42%/101%) | yes (27%/83%) | yes (31%/95%) |
| Flagellar basal-body rod protein FlgC | *flgC* | **BTB_RS08325** | **lmo0711_flgC** | **BSU16190_flgC** | yes (58%/100%) | yes (44%/103%) | yes (41%/103%) |
| flagellar hook-basal body protein FliE | *fliE* | **BTB_RS08330_fliE** | **lmo0712_fliE** | **BSU16200_fliE** | yes (52%/84%) | No | yes (26%/71%) |
| Flagellar MS-ring protein FliF | *fliF* | **BTB_RS08335** | **lmo0713_fliF** | **BSU16210_fliF** | yes (39%/97%) | yes (22%/99%) | yes (24%/77%) |
| Flagellar motor switch protein FliG | *fliG* | **BTB_RS08340** | **lmo0714_fliG** | **BSU16220_fliG** | yes (46%/95%) | yes (34%/99%) | yes (32%/96%) |
| Flagellar assembly protein H | *fliH* | **BTB_RS08345_fliH** | **lmo0715_fliH** | | yes (37%/104%) | no | no |
| Flagellum-specific ATP synthase FliI | *fliI* | **BTB_RS08350_fliI** | **lmo0716_fliI** | **BSU16240_fliI** | yes (70%/99%) | yes (44%/96%) | yes (44%/97%) |
| hypothetical protein | | **BTB_RS08355** | **lmo0694** | | yes (34%/99%) | no | no |
| hypothetical protein with C-terminal domain of FliK | | **BTB_RS08360** | **lmo0695** | | yes (34%/27%) | no | no |
| Flagellar basal body rod modification protein FlgD | *flgD* | **BTB_RS08365_flgD** | **lmo0696_flgD** | **BSU16280_flgD** | yes (45%/78%) | yes (31%/60%) | yes (33%/96%) |
| Fagellar hook-basal body protein, FlgE/F/G | *flgE/F/G* | **BTB_RS08370** | **lmo0697_flgE** | **BSU16290_flgG** | yes (47%/102%) | yes (37%/40%) | yes (39%/50%) |
| Protein of unknown function DUF3964 | | **BTB_RS08375** | **lmo0684** | | yes (41%/101%) | no | no |
| flagellar motor switch FliG superfamily protein | | **BTB_RS08380** | | | No | no | |
| Chemotaxis signal transduction protein CheV | *cheV* | **BTB_RS08385** | **lmo0689** | BSU14010_cheV | yes (64%/102%) | yes (49%/100%) | yes (49%/99%) |
| Motility gene repressor MogR | *mogR* | **BTB_RS08390** | **lmo0674_mogR** | | yes (35%/89%) | no | no |
| flagellin | *fla/hag* | **BTB_RS08395** | **lmo0690_flaA** | **BSU35360_hag** | yes (33%/104%) | yes (30%/109%) | yes (39%/105%) |
| flagellin | *flaA* | **BTB_RS08400** | | | no | no | |
| Putative lytic murein transglycosylase | | **BTB_RS08405** | **lmo0717** | BSU11570_yjbJ | yes (49%/82%) | yes (46%/69%) | yes (59%/56%) |
| Flagellar motor switch FliN | *fliN* | **BTB_RS08410** | **lmo0698** | | yes (50%/95%) | no | no |
| Flagellar motor switch protein FliM | *fliM* | **BTB_RS08415** | **lmo0699_fliM** | **BSU16310_fliM** | yes (49%/98%) | yes (26%/98%) | yes (25%/101%) |
| Flagellar motor switch FliN | *fliN* | **BTB_RS08420** | **lmo0693** | | yes (57%/70%) | no | no |
| Putative flagellar motor switch FliN | | **BTB_RS08425** | **lmo0675** | | yes (32%/97%) | no | no |
| Flagellar biosynthesis protein FliP | *fliP* | **BTB_RS08430** | **lmo0676_fliP** | **BSU16350_fliP** | yes (62%/84%) | yes (38%/93%) | yes (34%/89%) |
| Flagellar biosynthesis protein FliQ | *fliQ* | **BTB_RS08435** | **lmo0677_fliQ** | **BSU16360_fliQ** | yes (49%/98%) | yes (33%/93%) | yes (44%/60%) |

| | | | | | | | |
|---|---|---|---|---|---|---|---|
| Fflagellar biosynthesis protein FliR | *fliR* | BTB_RS08440 | lmo0678_fliR | BSU16370_fliR | yes (47%/100%) | yes (24%/98%) | yes (24%/93%) |
| Flagellar biosynthesis protein FlhB | *flhB* | BTB_RS08445 | lmo0679_flhB | BSU16380_flhB | yes (48%/100%) | yes (33%/97%) | yes (34%/98%) |
| Flagellar biosynthesis protein FlhA | *flhA* | BTB_RS08450 | lmo0680_flhA | BSU16390_flhA | yes (65%/100%) | yes (37%/100%) | yes (36%/101%) |
| Flagellar biosynthesis regulator FlhF | *flhF* | BTB_RS08455_flhF | lmo0681 | BSU16400_flhF | yes (33%/67%) | yes (29%/67%) | yes (34%/47%) |
| Flagellar basal-body rod protein FlgG | *flgG* | BTB_RS08460_flgG | lmo0682_flgG | | yes (42%/98%) | no | no |
| Flagellar motor protein MotA ($H^+$-coupled stator) | *motA* | BTB_RS22910 | lmo0685 | BSU13690_motA | yes (43%/96%) | yes (51%/99%) | yes (49%/91%) |
| Flagellar motor protein MotB ($H^+$-coupled stator) | *motB* | BTB_RS22905_motB | lmo0686_motB | BSU13680_motB | yes (45%/97%) | yes (48%/95%) | yes (44%/97%) |
| Glycosyltransferase. In *L. monocytogenes*: Flagellin glycosyltransferase and MogR antirepressor GmaR** | *gmaR**** | BTB_RS06215*** | lmo0688 | BSU21450_yolJ*** | yes (33%/42%) | yes (27%/54%) | yes (27%/53%) |
| hypothetical protein | | | lmo0701 | | no | | no |
| hypothetical protein | | | lmo0702 | | no | | no |
| hypothetical protein | | | lmo0704 | | no | | no |
| hypothetical protein | | | lmo0718 | | no | | no |
| Flagellar assembly protein H | *fliH* | | | BSU16230_fliH | | no | no |
| Flagellar export protein FliJ | *fliJ* | | | BSU16250_fliJ | | no | no |
| flagellar motor switch protein FliG superfamily protein | | | | BSU16260_ylxF | | no | no |
| Flagellar hook-length control protein FliK | *fliK* | | | BSU16270_fliK | | no | no |
| flagellar protein | | | | BSU16299_ylzI | | no | no |
| Flagellar basal body-associated protein FliL | *fliL* | | | BSU16300_fliL | | no | no |
| Flagellar biosynthesis protein, FliO | *fliO* | | | BSU16340_fliZ | | no | no |
| flagellum site-determining protein YlxH | | | | BSU16410_ylxH | | no | no |
| Chemotaxis protein, response regulator CheB | *cheB* | | | BSU16420_cheB | | no | no |
| Chemotaxis protein CheW | *cheW* | | | BSU16440_cheW | | no | no |
| CheY-P phosphatase CheC | *cheC* | | | BSU16450_cheC | | no | no |
| Chemoreceptor glutamine deamidase CheD | *cheD* | | | BSU16460_cheD | | no | no |
| RNA polymerase sigma-D factor | *sigD* | | | BSU16470_sigD | | no | no |
| swarming motility protein SwrB | *swrB* | | | BSU16480_ylxL | | no | no |
| Swarming motility protein SwrA | *swrA* | | | BSU35239-BSU35230_swrA**** | | no | no |
| ribosome hibernation promoting factor protein | *yvyD* | BTB_RS26450 | lmo2511 | BSU35310_yvyD | yes (63%/101%) | yes (67%/102%) | yes (60%/101%) |
| hypothetical protein | | | | BSU35319_yvzG | | no | no |
| flagellar protein FliT | *fliT* | | | BSU35320_fliT | | no | no |
| FlaG family protein YvyC | | | | BSU35350_yvyC | | no | no |
| Carbon storage regulator CsrA | *csrA* | | | BSU35370_csrA | | no | no |
| flagellar assembly factor FliW | *fliW* | | | BSU35380_yviF | | no | no |
| hypothetical protein | | | | BSU35390_yviE | | no | no |
| FlgN-like family protein | | | | BSU35420_yvyG | | no | no |
| negative regulator of flagellin synthesis | *flgM* | | | BSU35430_flgM | | no | no |
| Flagellar operon protein YvyF | *yvyF* | | | BSU35440_yvyF | | no | no |

* In parentheses: Percentage of identical matches in alignment / Percentage overlap of best hit, calculated as length of alignment relative to query protein length (both calculated as averages obtained from the forward and reciprocal blast)

** Score below threshold, bitscore 26.6 (87%/26%)

*** Although these proteins are reciprocal BLAST hits to *Lm* EGD-e GmaR, they are not considered true GmaR orthologs, as they lack the GmaR-specific C-terminal MogR-binding anti-repressor domain.

**** Laboratory strains of *B. subtilis* are known to contain an single “A” insertion mutation giving a frameshift in the *swrA* gene (McLoon et al., 2011). The protein used as query in this analysis was in-frame.

**Supplementary Table 2: Primers used in RT-qPCR**

| Gene name | Locus tag | Forward primer (5’-3’) | Reverse primer (5’-3’) | Product size | E | $r^2$ |
|---|---|---|---|---|---|---|
| *gatB* | BTB_RS21880 | agctggtcgtgaagaccttg | cggcataacagcagtcatca | 175 bp | 1.93 | 0.9968 |
| *rpsU* | BTB_RS21885 | aagatcggtttctaaaactggtaca | tttcttgccgcttcagattt | 102 bp | 1.87 | 0.9957 |
| udp | BTB_RS26495 | actagagaaacttggaaatgatcg | gacgcttaattgcacggaac | 101 bp | 1.85 | 0.9992 |
| *mogR* | BTB_RS08390 | gggatgcgagcatatgaaaa | aatgtttaaaccgtgacgttgac | 102 bp | 1.95 | 0.9992 |
| *flaB* | BTB_RS08395 | ctgcgaacggtacaaattca | aactcagtcgtctcgccaat | 100 bp | 1.96 | 0.9983 |
| *flaA* | BTB_RS08400 | ccgtgcaacactaggtgcta | cgtcttcgatttgagaagca | 104 bp | 1.94 | 0.9983 |

E, PCR efficiency. $r^2$, square of Pearson's correlation coefficient calculated from serial dilutions.

**Supplementary Table 3: Genome-wide gene expression analysis.** Differentially regulated genes identified by microarray-based transcriptional profiling of the MogR overexpression strain relative to an empty vector control. All genes with FDR-corrected *p*-values <0.05 are listed (ordered by locus tag number).

Genes downregulated in the MogR overexpression strain:

| Locus tag in *B. thuringiensis* 407 | Locus tag in *B. cereus* ATCC 14579 | Predicted function | $log_2$ (fold change) | *p*-value (FDR-corrected) |
|---|---|---|---|---|
| BTB_RS01475 | BC0294 | 10 kDa chaperonin | -1.06 | 0.00 |
| BTB_RS01480 | BC0295 | 60 kDa chaperonin | -0.58 | 0.05 |
| BTB_RS02175 | BC0422 | Methyl-accepting chemotaxis protein (with upstream "off" c-di-GMP riboswitch) | -1.31 | 0.00 |
| BTB_RS02990 | BC0587 | Acetyltransferase | -1.21 | 0.00 |
| BTB_RS03005 | BC0590 | Uncharacterized conserved protein YjgD | -0.73 | 0.03 |
| BTB_RS03040 | BC0598 | Helix-turn-helix domain protein/NprR | -0.89 | 0.00 |
| BTB_RS03260 | BC0643 | Amino-acid permease rocC | -0.63 | 0.02 |
| BTB_RS03410 | BC0666 | Immune inhibitor A | -1.03 | 0.00 |
| BTB_RS03430 | BC0670 | Phospholipase C (PC-PLC) | -2.32 | 0.00 |
| BTB_RS03435 | BC0671 | Sphingomyelinase C (Smase) | -1.47 | 0.01 |
| BTB_RS04285 | BC0791 | Coenzyme A disulfide reductase Cdr | -1.54 | 0.00 |
| BTB_RS04490 | BC0834 | Hypothetical protein | -1.95 | 0.00 |
| BTB_RS04760 | BC0888 | N-acetylmuramoyl-L-alanine amidase CwlH | -1.50 | 0.00 |
| BTB_RS05200 | BC1000 | Hypothetical protein | -1.01 | 0.00 |
| BTB_RS05215 | BC1003 | Serine-protein kinase rsbW | -0.91 | 0.00 |
| BTB_RS05220 | BC1004 | RNA polymerase sigma-B factor | -1.05 | 0.00 |
| BTB_RS05375 | BC1030 | Hypothetical protein | -0.51 | 0.05 |
| BTB_RS05580 | BC1061 | Hypothetical protein | -0.82 | 0.02 |
| BTB_RS05760 | BC1113 | Sigma-M negative effector | -1.61 | 0.01 |
| BTB_RS06105 | BC1185 | Dipeptide-binding protein DppE | -0.93 | 0.00 |
| BTB_RS06140 | BC1193 | Oligoendopeptidase F | -2.39 | 0.00 |
| BTB_RS06705 | BC1316 | Hypothetical protein | -0.66 | 0.03 |
| BTB_RS07240 | BC1424 | Ferredoxin--nitrite reductase NirA | -1.62 | 0.00 |
| BTB_RS07935 | BC1570 | Xanthine permease PbuX | -1.21 | 0.02 |
| BTB_RS08255 | BC1628 | Chemotaxis protein, histidine kinase CheA | -1.80 | 0.00 |
| BTB_RS08265 | BC1630 | Hypothetical protein | -0.92 | 0.03 |
| BTB_RS08295 | BC1636 | Flagellar hook-associated protein 1, FlgK | -1.67 | 0.02 |
| BTB_RS08310 | BC1639 | Flagellar (assembly) protein FliS | -0.68 | 0.00 |
| BTB_RS08325 | BC1642 | Flagellar basal-body rod protein FlgC | -2.87 | 0.00 |
| BTB_RS08335 | BC1644 | Flagellar MS-ring protein FliF | -2.31 | 0.00 |
| BTB_RS08340 | BC1645 | Flagellar motor switch protein FliG | -1.09 | 0.00 |
| BTB_RS08360 | BC1649 | Flagellar hook-length control protein FliK | -1.47 | 0.00 |
| BTB_RS08380 | BC1653 | Hypothetical protein | -1.37 | 0.00 |
| BTB_RS08385 | BC1654 | Chemotaxis signal transduction protein CheV | -1.80 | 0.00 |
| BTB_RS08395 | BC1656 | Flagellin FlaB | -1.04 | 0.00 |
| BTB_RS08400 | BC1657 | Flagellin FlaA | -3.45 | 0.00 |
| BTB_RS08405 | BC1660 | Putative lytic murein transglycosylase | -0.75 | 0.00 |
| BTB_RS08410 | BC1661 | Flagellar motor switch protein (fliN-homolog) | -0.99 | 0.00 |
| BTB_RS08430 | BC1665 | Flagellar biosynthesis protein FliP | -1.86 | 0.02 |
| BTB_RS08435 | BC1666 | Flagellar biosynthesis protein FliQ | -1.23 | 0.00 |
| BTB_RS08440 | BC1667 | Flagellar biosynthesis protein FliR | -1.02 | 0.00 |
| BTB_RS08445 | BC1668 | Flagellar biosynthesis protein FlhB | -1.60 | 0.00 |
| BTB_RS08450 | BC1669 | Flagellar biosynthesis protein FlhA | -1.43 | 0.00 |
| BTB_RS09080 | BC1789 | Transcriptional regulatory protein | -1.27 | 0.00 |
| BTB_RS10150 | BC1991 | Putative murein endopeptidase | -0.80 | 0.00 |
| BTB_RS10480 | BC2056 | Hypothetical protein | -0.74 | 0.02 |
| BTB_RS10590 | BC2076 | Acetyltransferase | -1.47 | 0.00 |
| BTB_RS12170 | BC2379 | Transcriptional regulator | -1.54 | 0.05 |
| BTB_RS12675 | BC2478 | ABC transporter, ATP-binding protein | -3.19 | 0.00 |
| BTB_RS12935 | BC2529 | Zn-dependent alcohol dehydrogenase | -1.97 | 0.00 |
| BTB_RS13545 | BC2659 | Glutamate-rich protein grpB | -1.72 | 0.00 |
| BTB_RS14470 | BC2832 | Putative aldehyde dehydrogenase DhaS | -2.10 | 0.00 |
| BTB_RS14520 | BC2842 | Hypothetical protein | -0.73 | 0.00 |
| BTB_RS14700 | BC2881 | Uncharacterized protein YhbF | -2.10 | 0.00 |
| BTB_RS14970 | BC2925 | Nucleotidyltransferase | -2.27 | 0.03 |
| BTB_RS15135 | BC2959 | Putative malate:quinone oxidoreductase Mqo | -0.67 | 0.00 |

| | | | | |
|---|---|---|---|---|
| BTB_RS15165 | BC2964 | Transcriptional regulator LsrR | -2.12 | 0.00 |
| BTB_RS12545 | BC3102 | Hemolysin BL-binding component HblA | -1.13 | 0.02 |
| BTB_RS12540 | BC3103 | Hbl component L1 | -1.59 | 0.00 |
| BTB_RS17315 | BC3461 | putative deacetylase YojG | -1.33 | 0.00 |
| BTB_RS17995 | BC3586 | Dipeptide-binding protein DppE | -1.15 | 0.00 |
| BTB_RS18825 | BC3762 | Serine protease, subtilase family Sfp | -0.84 | 0.02 |
| BTB_RS19450 | BC3835 | Ribonuclease HII | -1.24 | 0.00 |
| BTB_RS19615 | BC3867 | Putative coenzyme A biosynthesis bifunctional protein CoaBC | -1.12 | 0.00 |
| BTB_RS20130 | BC3966 | Hypothetical protein | -2.57 | 0.02 |
| BTB_RS20410 | BC4020 | Hypothetical protein | -0.98 | 0.02 |
| BTB_RS21125 | BC4154 | Hypothetical protein | -1.47 | 0.00 |
| BTB_RS21330 | BC4198 | Xaa-Pro dipeptidase PepQ | -0.51 | 0.01 |
| BTB_RS21660 | BC4260 | Glucokinase | -1.85 | 0.01 |
| BTB_RS21815 | BC4293 | Putative phosphotransferase | -0.71 | 0.04 |
| BTB_RS21885 | BC4307 | 30S ribosomal protein S21 | -0.81 | 0.00 |
| BTB_RS21915 | BC4313 | protein GrpE | -0.75 | 0.00 |
| BTB_RS21920 | BC4314 | Heat-inducible transcription repressor hrcA | -0.67 | 0.01 |
| BTB_RS22300 | BC4389 | Helicase, RecD/TraA | -1.53 | 0.00 |
| BTB_RS22905 | BC4512 | Flagellar motor protein MotB (H+-coupled stator) | -3.01 | 0.01 |
| BTB_RS22910 | BC4513 | Flagellar motor protein MotA (H+-coupled stator) | -1.74 | 0.00 |
| BTB_RS23315 | BC4594 | Citrate synthase 2 | -0.63 | 0.00 |
| BTB_RS23780 | BC4683 | putative ribosomal N-acetyltransferase YdaF | -1.06 | 0.00 |
| BTB_RS24165 | BC4756 | GDP-mannose-dependent alpha-mannosyltransferase MgtA | -1.58 | 0.00 |
| BTB_RS24190 | BC4762 | Phosphoenolpyruvate carboxykinase | -1.26 | 0.02 |
| BTB_RS22105 | BC4861 | hypothetical protein | -1.60 | 0.03 |
| BTB_RS25940 | BC5048 | Ferritin | -1.66 | 0.00 |
| BTB_RS26545 | BC5211 | putative permease IIC component YwbA | -1.23 | 0.02 |
| BTB_RS26650 | BC5228 | glycolate permease GlcA | -0.84 | 0.00 |
| BTB_RS26730 | BC5243 | hypothetical protein | -0.87 | 0.01 |
| BTB_RS26945 | BC5280 | 3R)-hydroxymyristoyl-[acyl-carrier-protein] dehydratase FabZ | -0.68 | 0.00 |
| BTB_RS27450 | BC5380 | putative siderophore-binding lipoprotein YfiY | -0.84 | 0.02 |

#### Genes upregulated in the MogR overexpression strain:

| **Locus tag in *B. thuringiensis* 407** | **Locus tag in Bc ATCC 14579** | **Predicted function** | **$log_2$ (fold change)** | **p-value** |
|---|---|---|---|---|
| BTB_RS02515 | BC0492 | Pyruvate formate-lyase-activating enzyme PflA | 0.92 | 0.00 |
| BTB_RS04390 | BC0813 | Cell wall-binding protein YocH | 0.96 | 0.00 |
| BTB_RS05575 | BC1060 | Collagen adhesion protein with upstream c-di-GMP "on" riboswitch CbpA | 3.02 | 0.00 |
| BTB_RS06050 | BC1174 | 3-oxoacyl-[acyl-carrier-protein] synthase 2 | 0.96 | 0.00 |
| BTB_RS06215 | BC1208 | Glycosyl transferase, group 2 | 1.23 | 0.00 |
| BTB_RS06545 | BC1283 | SinI | 1.08 | 0.05 |
| BTB_RS07850 | BC1553 | Penicillin-binding protein 1A/1B | 0.62 | 0.01 |
| BTB_RS08390 | BC1655 | MogR | 0.75 | 0.01 |
| BTB_RS09115 | BC1794 | Dipeptide-binding protein DppE | 2.01 | 0.00 |
| BTB_RS09415 | BC1853 | Hypothetical protein | 2.00 | 0.00 |
| BTB_RS10950 | BC2164 | Isoleucine--tRNA ligase IleS | 1.00 | 0.04 |
| BTB_RS11420 | BC2238 | Hypothetical protein | 1.95 | 0.00 |
| BTB_RS12700 | BC2483 | Putative acyl-CoA dehydrogenase YngJ | 1.81 | 0.00 |
| BTB_RS20160 | BC3972 | Pyruvate dehydrogenase complex E1 component, beta subunit | 0.71 | 0.01 |
| BTB_RS20730 | BC4083 | Putative adenine permease PurP | 0.73 | 0.03 |
| BTB_RS20740 | BC4085 | Pyrimidine-nucleoside phosphorylase Pdp | 0.80 | 0.02 |
| BTB_RS20745 | BC4086 | Purine nucleoside phosphorylase 1 | 0.87 | 0.01 |
| BTB_RS24745 | BC4870 | L-lactate dehydrogenase 2 | 0.66 | 0.02 |
| BTB_RS26190 | BC5138 | Phosphoglycerate kinase, Pkk | 0.83 | 0.00 |
| BTB_RS26200 | BC5141 | Central glycolytic genes regulator, CggR | 0.94 | 0.00 |
| BTB_RS26300 | BC5158 | Integral membrane protein | 1.58 | 0.02 |
| BTB_RS26705 | BC5238 | Glycine betaine transporter OpuD | 1.10 | 0.05 |
| BTB_RS26935 | BC5278 | Putative capsular polysaccharide biosynthesis protein YwqC | 0.71 | 0.02 |
